# Ecological Network Resilience & Extinction Proxies - Updating Projections of Ecological Networks

**DOI:** 10.1101/2023.08.02.551629

**Authors:** Erik Kusch, Alejandro Ordonez

**Author notes:** Corresponding Author: Erik Kusch.

## Abstract

Forecasting biodiversity and functioning changes to ecosystem composition and functioning under climate change requires using multi-species approaches that explicitly consider ecological interactions. Here, we propose a framework with which to incorporate considerations of (1) localised extinction risk proxies, (2) resilience mechanisms of ecological networks, and (3) extinction cascade directionality as a driving force of ecological change. These three aspects are seldomly considered when establishing ecosystems responses to climate change and biodiversity loss. Using this framework, we demonstrate that current practices may severely underpredict ecological change measured as loss of biodiversity and change in connectedness. Our novel framework which explicitly explores two-dimensional resilience landscapes defined by network resilience mechanisms (i.e., link loss sensitivity and realisation of rewiring potential) represents the most complete toolbox for assessment of vulnerability of ecological networks to extinction cascades. Ultimately, we propose that using localised extinction proxies, explicitly quantifying ecological network resilience through link-loss sensitivity and realisation of rewiring potential, as well as simulation of bidirectional extinction cascades will lead to improved capabilities of estimating ecosystem trajectories throughout the Anthropocene.

## Introduction

Ecological extinction events have been linked to ecosystem collapse^1^ and catastrophic consequences for human well-being^2^. This biodiversity loss crisis has been demonstrated to be driven predominantly by climate change and anthropogenic impacts such as land-use change^3–7^. Preventing ecosystem collapses and failures requires the identification of conservation needs according to projected losses of biodiversity in response to such extinction pressures throughout the anthropocene^8^.

Traditionally, assessing the risks to ecosystem processes and services exerted by climate change and other anthropogenic impacts has focused on establishing the net changes in biodiversity. This biodiversity accounting relies on modelling the distributions of individual species describing a species’ suitable habitat through its bioclimatic niche preferences^9^ and staking the suitability maps of many species^10^. However, this practice has been criticised as disregarding critical ecological processes such as interspecific interactions^11, 12^ or extinction cascades^13^. Central to the necessary refocusing towards community-level perspectives incorporating species-species interactions^14, 15^ is the analysis of ecological networks. These describe assemblages of species (i.e., nodes) and their interactions or associations (i.e., links between nodes)^16^. Incorporating such ecological networks into ecosystem projections (i.e., community composition and interactions between species) has been demonstrated to improve their accuracy, primarily through their capability of rendering extinction cascades (i.e., primary extinctions causing secondary extinctions), which have been identified as a critical driver of biodiversity loss at a global scale^17^. In this context, primary extinctions occur as critical thresholds in relevant species-specific extinction pressures are crossed leading to an initial set of extinct species. Secondary extinctions then occur as nodes lose links following primary extinctions thus becoming lesser connected or completely unconnected. This loss of links can be prevented by reallocation of lost links to extant nodes, which constitutes a rewiring of the original network^18, 19^.

Although efforts have been made to comprehensively include ecological networks in ecosystem projections, three knowledge gaps remain. First, species are subjected to several extinction pressures which establish primary extinction pools but there exists considerable ambiguity on how to quantify these pressures. Primary extinction risk proxies have been established through, among others: (1) range size, human forcing and habitat suitability^8, 20^, (2) projections of habitat loss^21^, (3) IUCN red list status^22–24^, (4) bioclimatic niche breadth^18^, and (5) climatic safety margins^25^. Among these, the IUCN red list status represents a global metric of extinction risk of a species. However, while this metric considers bioclimatic niches and projections of future climate envelopes, the global nature of this risk metric proves inaccurate at the local scales, where the impacts of biodiversity changes on ecosystem properties ought to be determined^26^. A localised metric of primary extinction risk are climate safety margins, which determine a species “typical” climatic limit^25, 27^. To establish scale-relevant proxies of extinction risk for network-driven ecosystem forecasts, the risk to ecosystem integrity needs to be compared when using global and local primary extinction risk proxies.

Second, the interaction between primary and secondary extinctions determine network resilience to primary extinctions, which can be defined as the balance between the resistance (i.e., sensitivity to link loss)^18^ and recovery (i.e., the realisation of rewiring potential)^28, 29^ of species interactions (i.e., links). By integrating link-loss resistance and recovery, we may formulate a two-dimensional resilience landscape in which the likely consequences of primary extinctions can be explored for any combination of resistance and recovery potential of any given ecological network (see Figure 1). Studies of the impacts of biodiversity changes in ecological networks largely fail to integrate either of these critical mechanisms fully. For instance, network links are commonly treated as binary states (i.e., present or absent) despite there being reasons to believe that species may experience significant extinction pressure through the loss of particularly strong connections rather than the number of connections themselves^30^. As an example, loosing access to a singular food item constituting 50% of one’s diet may be much more detrimental than loosing access to 50% of food items so long as their combined proportion in the total diet does not exceed 50%. As such, link-loss sensitivity can be understood as a percentage-based threshold below (e.g., a link-loss sensitivity of 0.6 would indicate secondary extinction when a given node retains less than 60% of its initial combined link weight due to removal of other nodes from the network). Additionally, ecological networks are usually considered as static assemblages without any possibility of rewiring following a compositional change. Ultimately, most studies of extinction cascades have only explored one specific location across network-resilience landscapes (i.e., location IV in Figure 1C) instead of generating a range of ecologically feasible ecosystem projections. Although some studies have begun to explore the network resilience landscapes^18^, we must quantify potential future scenarios of ecosystem projections within sensible ranges across network resilience landscapes. Doing this would provide a deeper understanding of how link-loss sensitivity and the realisation of rewiring potential could affect future biodiversity levels.

**FIGURE 1.**
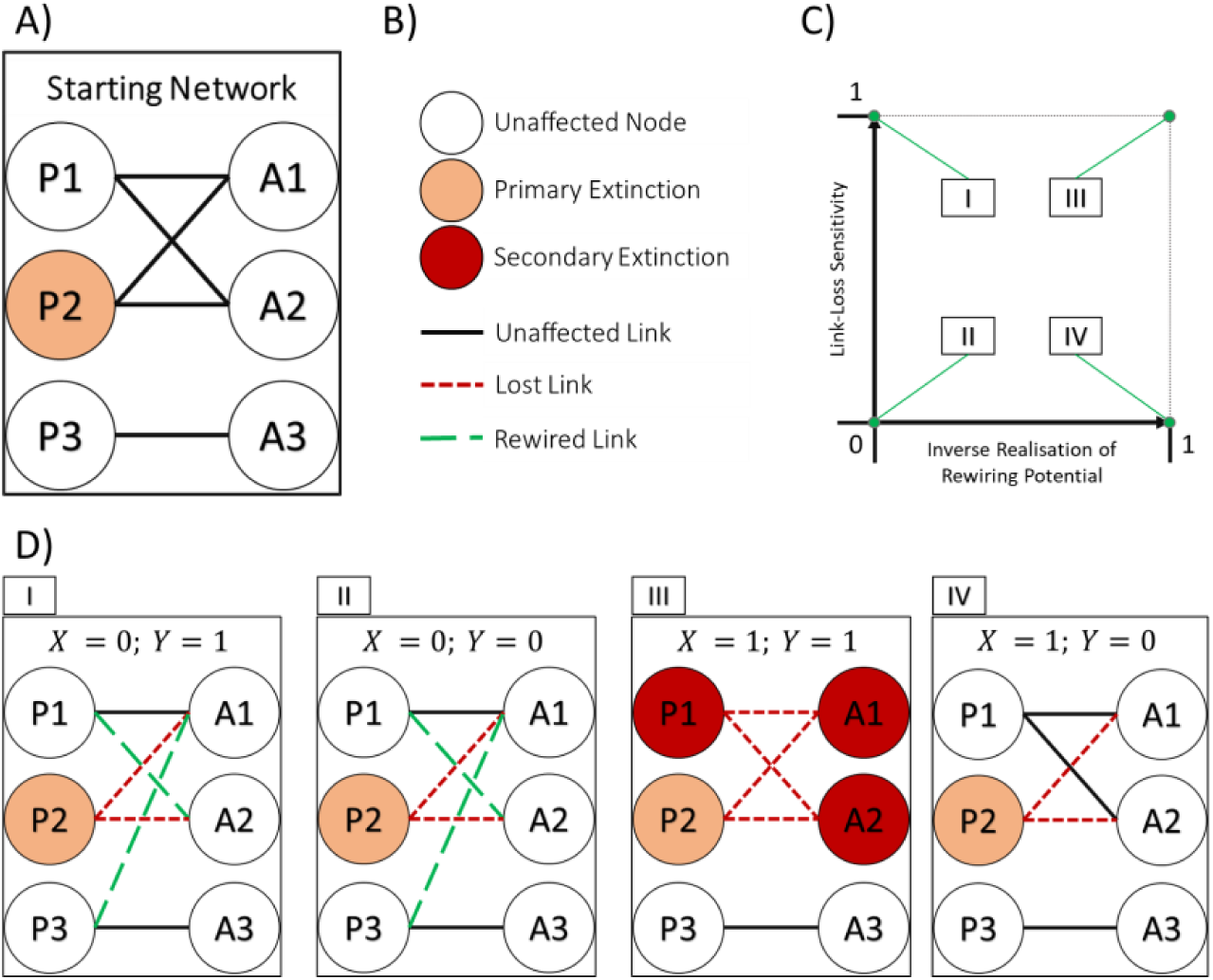
THE NETWORK RESILIENCE LANDSCAPE CONCEPT. Using the concept of a resilience landscape as depicted in C and defined by link-loss sensitivity (Y axis) and inverse realisation of rewiring potential (X axis), it is possible to explore post-extinction cascade scenarios of any original network (an example is shown in A) according to resistance and recovery mechanisms in response to primary extinctions (node P2 in network visualisations) according to X and Y coordinates across the network resilience landscape. Examples of such network scenarios are shown in D. Refer to B for a legend of network visualisation items.

Lastly, extinction cascades in multitrophic networks can be categorised for most ecological networks either as bottom-up or top-down processes. Bottom-up extinction cascades are characterised by the primary removal of basal species forcing secondary extinctions of higher-order species^17^. Top-down extinction cascades, on the other hand, highlight how the removal of higher-order species can result in drastic changes in basal species’ population dynamics, which may disrupt ecosystem processes sufficiently to trigger secondary extinctions^31^. A contemporary analysis of bottom-up and top-down extinction severity in mutualistic networks highlighted that biodiversity changes are larger following bottom-up extinctions than top-down extinctions^18^. However, these results may not be generalisable, making it pivotal to quantify the relative contributions of top-down and bottom-up extinctions as multipliers of environmental driven biodiversity changes.

Leveraging a global collection of plant-frugivory networks^32^, we first quantify global and local extinction risk proxies to compare primary extinction classifications (i.e., species identities which are under primary extinction risk) and extinction simulation outcomes (i.e., ecological networks after primary and secondary extinctions have been accounted for). We expect local extinction risk proxies to identify larger pools of primary extinctions due to the effect of refugia preventing global extinction of a species irrespective of local conditions in any specific network location^33^. Second, we quantify the change in species compositions and ecological network topological metrics respective to positioning within network-resilience landscapes, thus exploring potential scenarios of ecosystem change. We propose that the severity of extinction cascades decreases with link-loss sensitivity and increases in the likelihood of the realisation of rewiring potential. Such a gradient would indicate that current practices of extinction cascade predictions may underpredict biodiversity loss due to their simplistic implementation of either network resilience mechanism. Third, we contrast the effects of bottom-up and top-down extinction cascade effects to build a better understanding of extinction cascade risk-mitigation. In line with previous research, we expect mutualistic networks to be more affected by bottom-up than top-down extinction cascades^18^, but suggest that bidirectional extinction cascades will lead to considerably greater biodiversity loss than unidirectional cascade scenarios. As such, this study represents an investigation of future ecological network and ecosystem scenarios which is unparalleled in its completeness and integration of key concepts of secondary extinction research.

## Material & Methods

All data handling and analyses were carried out in the R environment^34^. For a visual representation of the data handling, extinction proxy identification, rewiring probability quantification, and extinction simulation workflow see Figures S1, S2 and S3, respectively.

### Network Data

We obtained ecological network data from Fricke, 2021^35^, who compiled information on 406 plant-frugivory networks sampled in-situ^35^. The quantification of links between nodes varies among these networks. While some networks report link strength as the number of fruits consumed^36^, other networks store interaction strength as a combination of fruit-handling time and behaviour^37^. Nevertheless, due to homogenous units of link-weight within each individual network, the impact of link-loss sensitivity (a percentage-based threshold) can be compared across all networks. To study the effects of varying link-loss sensitivity, we select from this pool of 406 ecological networks only those who display a considerable variation in the strength of their realised (i.e., non-zero) interactions. We quantify this variation as the coefficient of variation of realised link strengths and retain only those networks for which this value exceeds 0.5 (i.e., the standard deviation of interaction strengths must be at least half as large as their mean). In addition, to ensure that the networks in our analysis represent complex ecosystems with potential extinction cascades, we exclude all ecological networks composed of less than seven frugivores and seven plant species. Following these criteria, our analyses focus on 81 networks spanning all continents except Antarctica (see Figure S4 for network observation locations).

### Climate Data Retrieval

To identify primary extinction orders, we obtained air temperature and soil moisture for the locations of all 81 study networks. We did so for the 1982-1999 period (i.e., historical baseline) and 2081-2100 (i.e., climate projection). We focus on establishing extinction proxies through these two essential climate variables since temperature has been linked to many fundamental physiological processes in animals^38^ and plants^39^ while soil moisture has been shown to determine vegetation properties^40^. For the historical baseline, we obtained data from ERA5-Land climate reanalysis^41^ using the KrigR R-package^42^. For future conditions, we obtained data from the Coupled Model Inter-comparison Project Phase 6 (CMIP6)^43^ full Earth System Model simulations of historical climate and projections for the end of the 21^st^ century (i.e., 2081-2100). We selected the ssp245 future scenario, which represents the “middle of the road” climate commitment with climate protection measures being taken. In addition, we also selected the ssp585 scenario to explore ecological extinction cascades under the “worst-case” emission scenario. Using KrigR, we statistically downscaled CMIP6 to a 9×9km resolution aligning with the native ERA5-Land raster. This procedure has previously been demonstrated to yield accurate and robust outputs for our two target variables^44^. To reduce the computational cost of these downscaling efforts, we kriged raster products in a 5° buffer around the location of each study network rather than at a global extent. We subsequently created downscaled, bias-corrected climate projections by summing historical ERA5-Land baseline data with the difference between the two downscaled CMIP6 products following previously established practice^42, 45–47^.

### Species Occurrence Data

The 81 networks we retained for our analyses contained 1916 species (1123 plant species, 793 animal species). To represent species distributions for the calculation of extinction proxies, we obtained occurrence records for each species from GBIF^48,49^ using the rgbif package^50^. We limit species occurrence retrieval to the same period as the historical baseline climate data, ensuring the occurrence information can be used to describe the sampled species climatesafety-margins^25^. Species with less than 20 occurrence records we removed from our analyses. This resulted in the removal of 192 species (155 plant and 37 animal species). To remove potentially erroneous species occurrence records from the data we retrieved, we used open-source shapefiles^51^ to establish centroid locations for continents, countries, and states. Subsequently, we removed occurrence records with a distance to the centroid of a location larger than .001° (∼111m) which did not reduce the number of occurrences per species to below 20. Furthermore, we removed those observations with climatic conditions larger than one standard deviation around the species-specific mean to avoid biasing climate safety margin computation with unrepresentatively wide climate tolerances.

### Extinction proxies

We use three different extinction proxies (i.e., IUCN, climate-safety-margins, and node-centrality) to carry out primary extinctions across the 81 evaluated networks.

#### IUCN

For each species, we queried IUCN red list classification from IUCN records using the rredlist R package^52^. Doing so, we retrieved IUCN red list classifications for 1244 of our 1724 species. For the remaining 480 species, we used the ConR R package^53^ to calculate IUCN red list status. To do so, the ConR package estimates extent of occurrence (EOO), area of occupancy (AOO) and an estimate of the number of locations. To account for the effect of protected areas on IUCN status in the ConR estimation, we supplied the algorithm with shapefiles from the World Database on Protected Areas (WDPA)^54^.

#### Climate-safety-margins

We used the climate-safety-margin concept (a “typical” species observed climatic limit^25^) adapted for a single-species design. With this approach, we can localise extinction risk per species for each network location rather than using global classifications as supplied by IUCN red list status. We first identify climatic preferences for each species. To do so, we extracted values corresponding to air temperature and soil moisture experienced by each species based on the historical baseline climatic conditions defined by their occurrences. We then established the median and standard deviation of bioclimatic niche realisation per species to define a “bioclimatic profile”. Next, we identified future projections of climatic conditions at the location of each ecological network in our analysis using our bias-corrected, downscaled end of the 21^st^-century climate projections (ssp245 and spp585). Lastly, we quantified the localised climate-change-driven extinction risk (C*_i,j_*) for each species (*i*) at each network location (*j*) as the absolute difference between climate preference (*M_i_*, median of bioclimatic niche) and projected conditions at each network’s location (*P_j_*) divided by the bioclimatic niche breadth (*σ_i_*, standard deviation of the bioclimatic niche):

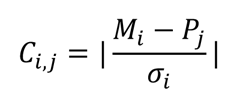

#### Centrality

To establish a worst-case scenario of biodiversity loss, we also simulate extinction cascades driven by the primary extirpation of the most central species in each network (i.e., the most connected species). The centrality of a node has been demonstrated to be a driver of secondary extinctions, with the removal of more central nodes leading to more secondary extinctions^55^. We identify the most connected species for each network by assessing node strength (i.e., the sum of all link weights connected to each node^56^). It is essential to clarify that we use this method of establishing a primary extinction sequence as a worst-case baseline for each network instead of as an attempt to predict future ecosystems accurately.

### Rewiring Potential

A database linking functional group membership and functional trait expression of plants and frugivores to binary interaction outcome (whether a frugivore consumes fruits of a plant) is also part of Fricke, 2021^35^. We use this database to establish the potential of each frugivore to interact with each plant and vice-versa (i.e., the potential of a plant’s fruit being consumed by each frugivore). We do so with the randomForest R package^57^ using the entire database (including networks not selected for our analysis presented here). To infer the probability of a frugivore species consuming the fruit of each given species, we informed the random forest algorithm with data about plant vegetative height, seed dry mass, seed length, seed thickness, seed width, specific leaf area, stem-specific density, and fruit length. To infer link-probability for each plant with each frugivore species, we used functional group membership, mean body mass, and diet preferences of each frugivore as predictors in our random forest models. Predictions of rewiring probability were made on a network-by-network basis using only functional expressions of species belonging to each network to generate species-and network-specific matrices of the probability of interactions occurring between frugivore-plant pairs.

### Extinction Simulation

Simulations of extinctions and subsequent secondary extinctions were carried out using functionality of the NetworkExtinction R package^19^, which exports post-extinction simulation adjacency matrices and enables full integration of network resilience mechanisms as well as full tracking of extinction cascades (i.e., beyond first-order secondary extinctions).

To study primary extinctions and subsequent secondary extinctions determined by different extinction proxies, we selected four sets of primary extinction species pools per network. Using IUCN red list status, we selected for primary extinction all species whose status was either endangered or critically endangered. For the Climate-safety-margins proxy, a species was removed if future projected temperature or soil moisture at a network location is at least two standard deviations removed from the current bioclimatic niche median for a species (i.e., extinction proxy exceeds 2) were selected for primary extinction under each emissions scenario respectively (i.e., ssp245 and ssp585). Lastly, to establish a worst-case baseline using centrality scores, we identified the top 0.25 percentile of species regarding node strength per network as primarily extinct species. For climate-safety-margin-and centrality-driven primary extinction identification, order of primary extinction was defined by magnitude of extinction proxy (e.g., higher climate-safety-margin proxy species go extinct before lower climate-safety-margin extinction proxy species). For IUCN-driven extinction analyses, species were removed in alphanumerical order, within each status category with critically endangered species always being removed before endangered species. For each set of primary extinction species pools, we ran one simulation of extinction cascades per network (4 extinction proxies * 81 networks = 324 simulations).

To implement and quantify the impacts of ecological network resilience mechanisms on ecosystem projections, we utilised the functionality of the NetworkExtinction package to simulate extinction cascades at varying levels of link-loss sensitivity and rewiring potential realisation. First, establishing link-loss sensitivity, we identified secondary extinctions as all species that retain less than a given percentage of initial interaction strength because of the removal of adjacent nodes in their network. Secondly, in realising rewiring potential, we used rewiring probabilities calculated for each species-pairing in each network and simulated instantaneous rewiring of a link lost due to the removal of a node in an interaction pair. This rewiring was only allowed if a given rewiring probability threshold was exceeded and rewiring happened only to the most likely partner. For example, a species losing a link was permitted to add the strength of the said link to the strength of another link (even if such a link is an unrealised connection) if the probability of the species that lost their original partner rewiring to their novel partner exceeds the rewiring probability threshold. We execute simulations of all combinations of link-loss and rewiring probability thresholds ranging from 0 to 1 in increments of 0.05 (21 increments), effectively raising the number of total simulations to 142,884 (324 simulations * 21 link-loss thresholds * 21 rewiring probability thresholds). At their extremes, these values are interpreted as follows. A secondary extinction threshold of 0 signifies secondary extinctions manifesting only when a node becomes completely unconnected. In contrast, a value of 1 means that a node may not lose any interaction strength to withstand becoming secondarily extinct. A rewiring probability threshold of 0 indicates that a species may rewire all its connections to any species for whom the rewiring probability is greater than 0 (i.e., virtually all other nodes of the same frugivory-plant grouping). Meanwhile, a value of 1 implies that rewiring may only happen to partners with a rewiring probability of 1 (i.e., virtually no other node). Exploring the ecosystem projections generated across the combinations of these two thresholds allows us to explore an array of optimistic, pessimistic, and realistic biodiversity projections.

To identify the impact of extinction cascade directions, we ran our simulations using different primary extinction species pools. To establish the most realistic projections of ecosystems, we used the above-defined primary extinction species pools as-is. These represent extinction cascades starting at the producer and consumer level (i.e., bidirectional cascades). To quantify the effects of bottom-up and top-down extinction cascades we only retained plant and animal species within the primary extinction species pool, respectively. For computational reasons we executed only bidirectional cascade simulations for the ssp585 scenario. Consequently, the total number of extinction simulations rose from to 357,210.

Lastly, to identify how much given extinction proxy and cascade direction change ecosystems (as measured through loss of biodiversity and change in network topology) deterministically as opposed to random species removals, we execute 100 simulations for random primary extinction species pools with the same number of species as selected by each extinction proxy for each extinction cascade direction thus bringing the final total of simulations run for this study to 36,078,210 (357,210 deterministic simulations, 35,721,000 random simulations).

### Analyses

To explore the impacts of choice of primary extinction risk proxy, network resilience landscape placement and extinction cascade directionality, we calculated a set of two ecosystem change metrics for each simulation outcome. The first summarises relative biodiversity change relative to pre-extinction biodiversity levels for quantifying relative species loss. The second, quantifies changes in overall ecosystem structure through a relative change metric of network topology by quantifying relative link loss. To quantify the implications of primary extinction proxies and extinction cascade directionality on our two relative ecosystem change metrics, we executed Bayesian models using the brms R package^58^. Using brms, we established models for each ecosystem change metric as explained through additive effects of link-loss and rewiring probability thresholds and their interaction. To analyse the effect of different primary extinction proxies, we restricted our model data to bidirectional simulation runs and added an additive coefficient of the extinction-risk proxy being used. To quantify the effect of extinction cascade directionality, we restricted our model data to outcomes driven by the ssp245 climate-safety-margin proxy and added an additive coefficient of cascade directionality to the model. Our models used a zero-one inflated beta distribution for their outcome variables, as their values must fall between 0 and 1.

## Results

### Extinction Proxy Identification

Across all 81 networks, the global IUCN classifications identify seven species as going primarily extinct (*Ateles belzebuth, Chloropsis cochinchinensis, Coffea arabica, Loxodonta africana, Myrcianthes pungens, Tectona grandis, Vitex cooperi*). This species list corresponds to approximately 0.41% of our 1724 species. By comparison, ssp245 climate-safety-margins selected 595 unique species identities for primary local extinction which is almost doubled under ssp585 with 1031 unique species in the primary local extinction pools. Our worst-case baseline of removing the top 25% quantile of most central species per network identifies a similar number of species, but not entirely same sets of primary extinction species as the ssp245 climate-safety-margin selection procedure (506 unique species identities, 187 of which are shared between the two localised extinction risk proxies). See Figure S5 for a visual representation of these primary extinction species pools and their overlaps. Our comparison highlights that global assessments of primary extinction risk are likely to underpredict primary extinction risks by identifying too few primary extinctions than what may be realistic at local scales.

### Network-Resilience Landscapes

To begin, we focus on biodiversity loss within a Peruvian plant-bat mutualistic frugivory network^59^ at different positions across the network-resilience landscape (Figure 2) while taking into account link-loss sensitivity (Y-axis in Figure 2B) and rewiring capabilities abating secondary extinction (X-axis in Figure 2B). Note that each location across the network-resilience landscape in Figure 2 represents a distinct ecosystem projection driven by the same ssp245 climate-safety-margin driven set of primary extinctions following a bidirectional extinction cascade. Depending on position across the network-resilience landscape, ecosystem projections resulting from the same primary extinctions change drastically with increased link-loss and rewiring probability thresholds leading to a greater relative loss of species, particularly when increasing simultaneously. In the most extreme case (position 9 in Figure 2), where rewiring is impossible and any loss of interaction strength leads to secondary extinction, all nodes (i.e., species) connected to the primary extinction set either directly or indirectly are lost. At the other extreme of rewiring probability thresholds, however, we find that completely free rewiring of any lost network connection can halt the network level effects of secondary extinctions irrespective of link-loss thresholds (positions 1, 2 and 3 in Figure 2).

**FIGURE 2.**
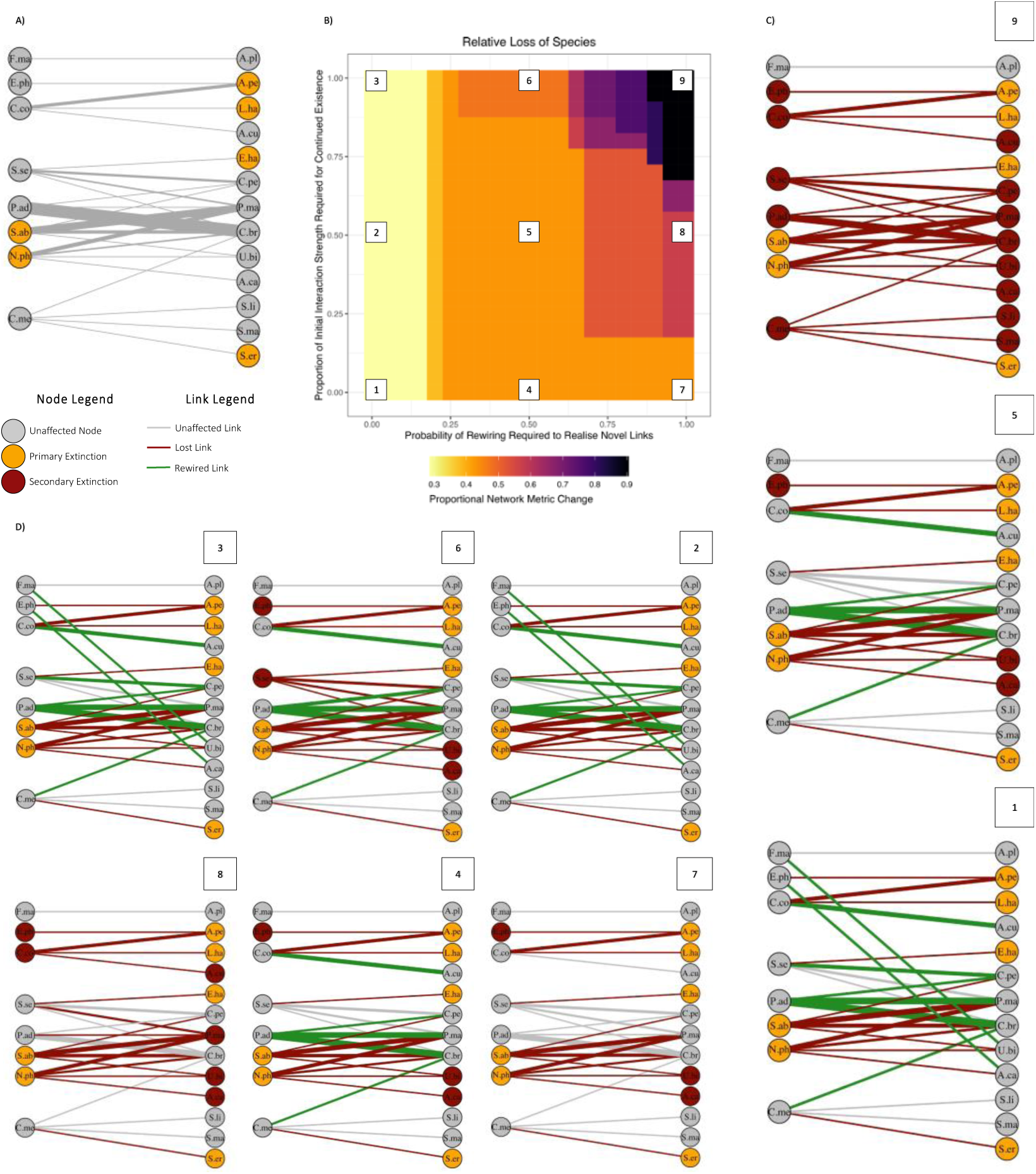
ECOLOGICAL NETWORK RESILIENCE MECHANISMS DETERMINE POST-EXTINCTION CASCADE ECOSYSTEM CONSTELLATIONS. Example outcome of extinction simulation driven by climate safety margin extinction proxy for example plant-frugivory network obtained from Arias Arone, 2016^59^ and contained within Fricke, 2021^35^ (A). Network-resilience landscape of relative species loss (B) shows that increased link-loss and increased rewiring probability thresholds lead to greater relative loss of species particularly when increasing simultaneously. Individual ecosystem projections located across the network-resilience landscape are shown in C) and D). Three-letter codes in nodes refer to species identities: A.ca … *Anoura caudifer*, A.cu … *Anoura cultrata*, A.pe … *Anoura peruana*, A.pl … *Artibeus planirostris*, C.br … *Carollia brevicauda*, C.co … *Condaminea corymbosa*, C.me … *Cecropia membranacea*, C.pe … *Carollia perspicillata*, E.ha … *Enchisthenes hartii*, E.ph … *Epiphyllum phyllanthus*, F.ma … *Ficus maxima*, L.ha … *Lonchophylla handleyi*, N.ph … *Nicandra physalodes*, P.ad … *Piper aduncum*, P.ma … *Platyrrhinus masu*, S.ab … *Solanum abitaguense*, S.er … *Sturnira erythromos*, S.li … *Sturnira lilium*, S.ma … *Sturnira magna*, S.se … *Solanum sessile*, U.bi … *Uroderma bilobatum*. Link width indicates logarithmic interaction strength but is not consistent between network plots.

Analysing the mean of ecosystem change metrics for all 81 evaluated networks, we find a generalized change in the ecosystem and network metrics due to shifts in link-loss and an increase in rewiring probability (Figure 3). Increasing link-loss and rewiring probability thresholds results in a larger change in ecosystem summary metrics irrespective of metrics of change (Figure 3, 4, and 5), primary-extinction-defining proxies (Figures S6, S7, and S8) or cascade directionality (Figures S10 and S11). Finally, we established effect sizes for each simulation outcome across the network-resilience landscape for all extinction risk proxies when assessing bidirectional extinction cascades. We used these metrics to contrast deterministic simulation outcomes with randomly selected primary extinction sets of the same number of species as the deterministic counterparts. These contrasts show no generalisation for effect sizes of IUCN or climate-safety-margin derived primary extinction sequences (Figure S9). What these contrasts do show is that, as expected, removing species by their network-specific centrality scores leads to much more drastic losses of biodiversity and network links than their random simulation counterparts.

**FIGURE 3.**
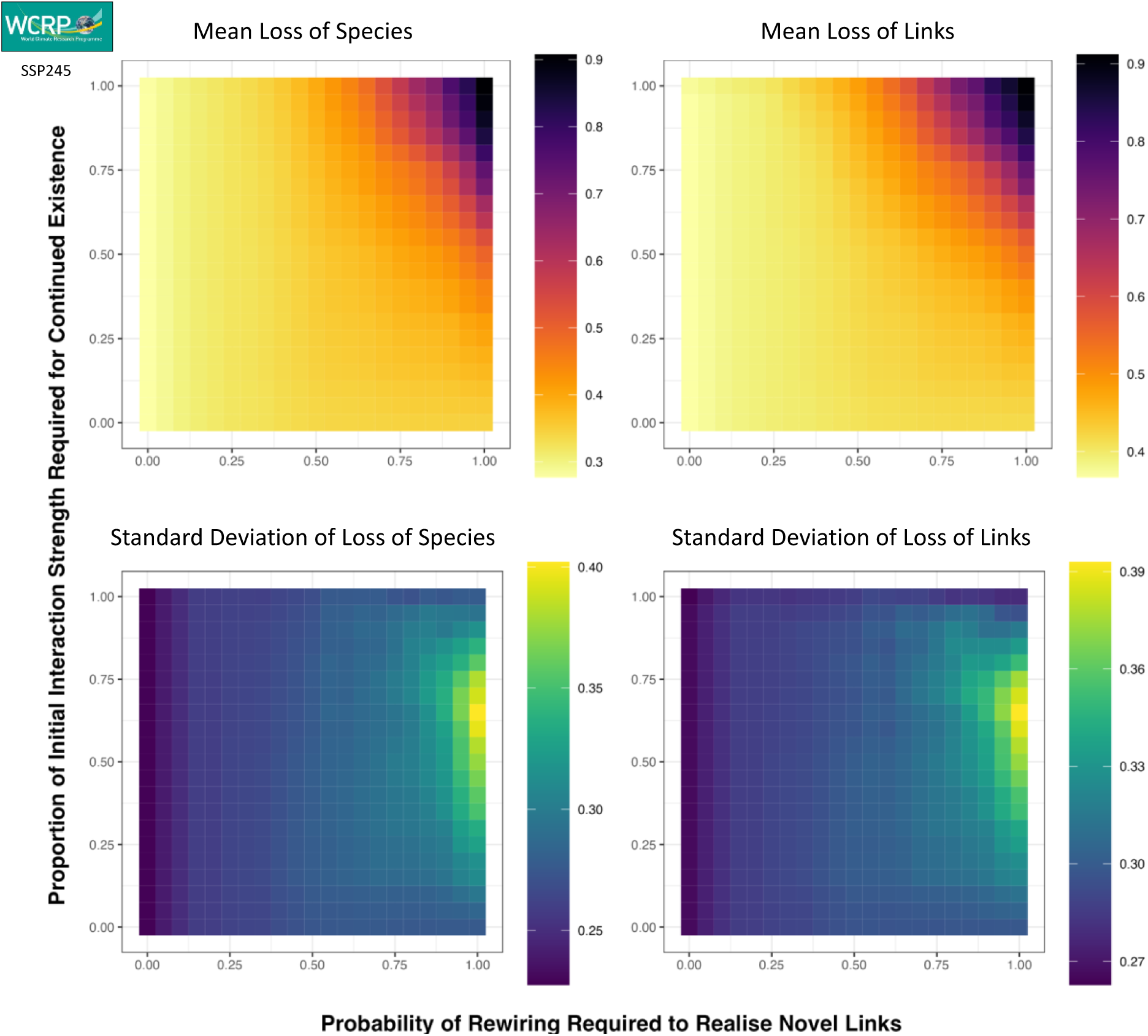
NETWORK-RESILIENCE LANDSCAPES ARE GENERALISABLE IRRESPECTIVE OF TARGET METRIC. Mean network-specific relative change in number of species, number of interactions/links, for all 81 study networks demonstrates the generality of the network-resilience landscape concept as pictured for a single network in Figure 2B. Note that these results show the extinction simulation outcomes of bidirectional extinction cascades as driven by primary extinctions characterised through ssp245 climate-safety-margin proxies. For comparable visualisations of bidirectional cascades driven by IUCN, Centrality and ssp585 climate-safety-margins, please refer to Figures S6, S7, and S8, respectively.

### Drivers of Ecosystem Changes

When quantifying the magnitude of ecosystem and network change owing to extinction cascade dynamics, we investigate how extinction proxies affect our target metrics of change (relative loss of nodes and links per network) following bidirectional extinction cascades while we investigate the effects of extinction cascade directionality using ssp245 climate-safety-margin informed simulations.

#### Extinction Risk Proxies

To identify the change in network and ecosystem constellations owing to differences in primary extinction species pools of different extinction proxies we evaluate relative coefficient magnitudes of our ecosystem change metric (i.e., biodiversity and link loss) models. Figure 4 shows differences in relative species loss and link loss between extinction proxies. Primary extinctions driven by node centrality led to the greatest changes in both target metrics, followed by those informed through ssp585 climate-safety-margins and ssp245 climate-safety-margins. Notably, extinction cascade outcomes, particularly in remark to biodiversity loss, which are to be expected under the worst-case emission scenario ssp585 are very similar to those under our “worst-case” centrality removal baseline (see for this also Figures S7 and S8). Ultimately, we find that extinction-cascade-driven ecosystem change is much more pronounced when using localised proxies rather than global ones.

**FIGURE 4.**
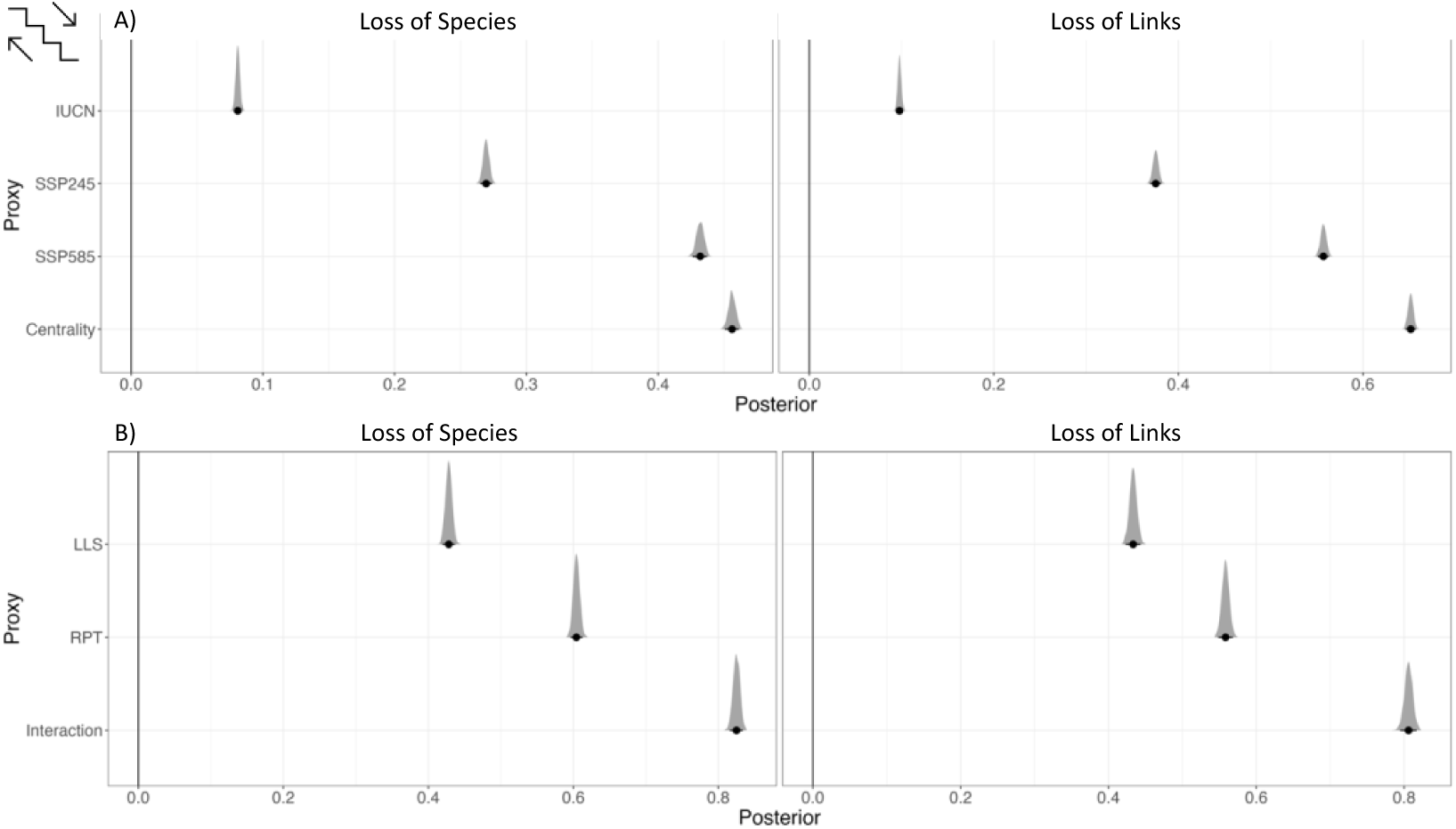
LOCALISED EXTINCTION RISK PROXIES LEAD TO GREATER CHANGE IN NETWORK AND ECOSYSTEM METRICS THAN GLOBAL EXTINCTION RISK PROXIES. A) Primary extinction sets defined through node centrality led to greatest changes in ecosystem and network metrics while localised climate-safety-margin determined primary extinction sequences lead to intermediate changes with IUCN-informed primary extinctions spurring the least change in networks through extinction cascades. Note, however, the similarity of biodiversity loss under ssp585 and at the “worst-case” centrality-removal case. This observation holds true for all target ecosystem and network metric we analyse here. Icon on the top-left identifies bidirectional cascade simulations in line with visualisations in Figure 5 and supplementary figures. B) Increases in resilience process thresholds (i.e., link loss sensitivity – LLS; rewiring probability threshold – RPT; and their interaction) lead to increases in species and link-loss.

#### Cascade Directions

When quantifying the effects of extinction cascade directionality on ecosystem changes, we focus on extinction simulations involving primary extinction sequences determined through ssp245 climate-safety-margins. We find that, irrespective of network/ecosystem change metric, bidirectional extinction cascades lead to the largest change in ecosystems while bottom-up cascades lead to a greater relative loss of species and links than top-down extinction cascades (Figure 5). This holds true for all primary extinction proxies for which all cascade directionalities were simulated (i.e., ssp245 climate-safety-margins, IUCN, and centrality; see Figures S10 and S11.

**FIGURE 5.**
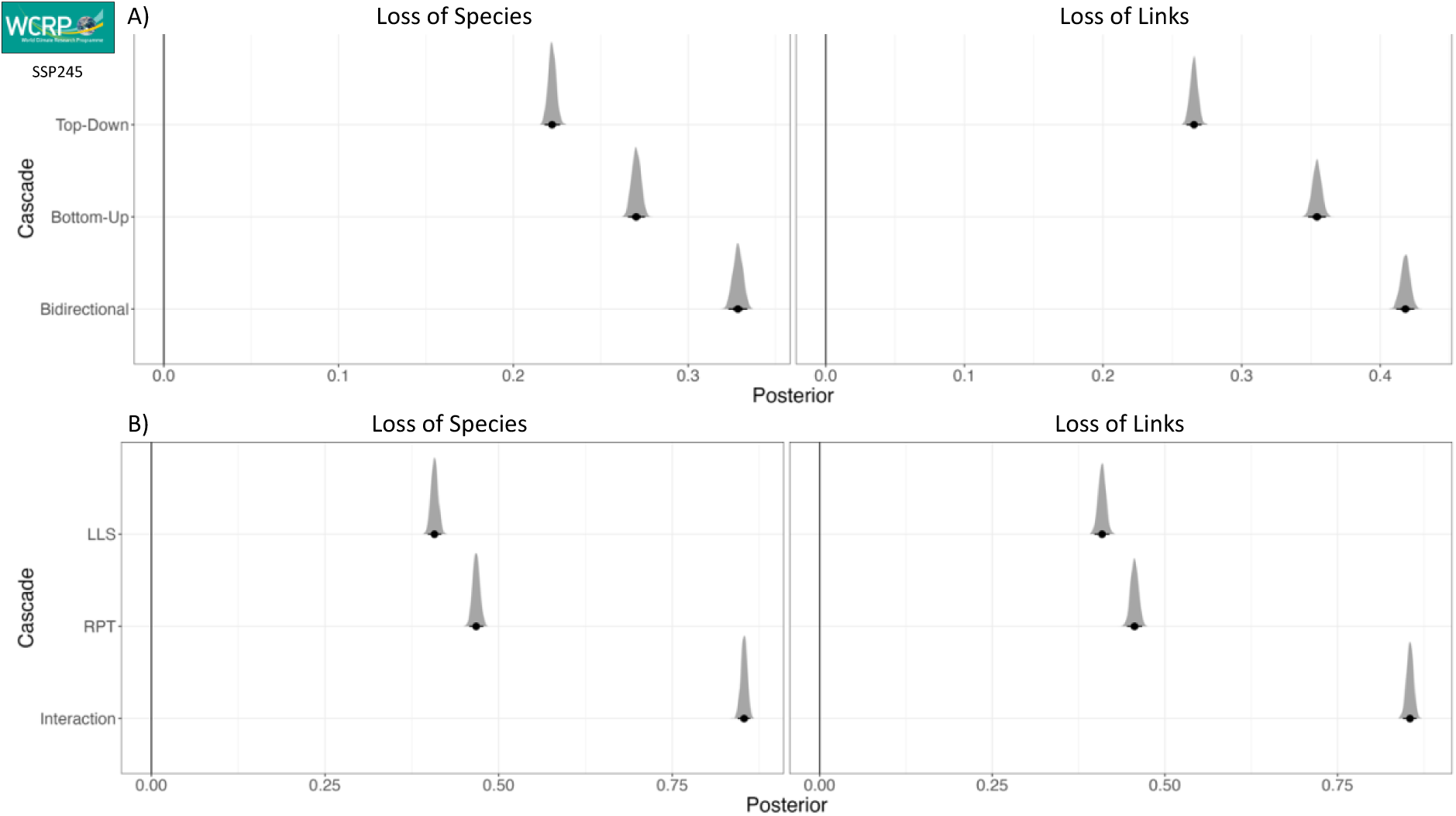
BIDIRECTIONAL EXTINCTION CASCADES LEAD TO GREATER CHANGE IN ECOSYSTEMS THAN TOP-DOWN OR BOTTOM-UP CASCADES. As our simulations show, both bottom-up and top-down cascades affect changes in ecosystem and network metrics when contrasting pre-and post-extinction simulation networks. While top-down cascades overall lead to smaller changes than bottom-up cascades, both are outdone in change magnitude by bidirectional extinction cascades. See Figures S10 and S11 for the overview of differences in network-landscape representation. B) Increases in resilience process thresholds (i.e., link loss sensitivity – LLS; rewiring probability threshold – RPT; and their interaction) lead to increases in species and link-loss.

## Discussion

Our study identifies an extensive range of ecologically feasible ecosystem projections which highlight how extinction cascade forecast practices shape our understanding of ecosystem trajectories in the face of relevant extinction pressures throughout the Anthropocene. The three key knowledge gaps we aim to fill with this study showed substantial implications for conservation that need to be addressed through ecological networks.

### Extinction Risk Proxies

Our overview of primary extinction sequences as identified through IUCN red list status and localised climate-safety-margin extinction risk proxies yield vastly different understandings of how many species are projected to go extinct thus potentially leading to secondary extinctions. We find that, according to our chosen thresholds and assuming agreed-upon climate change mitigation action is taken at a global level (i.e., ssp245), global extinction risk proxies identify 85 times fewer primary extinctions than localised extinction risk proxies, with a minimal agreement in species identity among primary extinction sequences between the two (see Figure S5). However, current trajectories of emission scenarios place future projections of environmental change closer to the ssp585 scenario under which global extinction risk underpredicts primary extinctions species identities by a factor of 147. This confirms our expectation of global extinction risk proxies inferring less extinction events relevant to studies of extinction cascades than do localised proxies of extinction risk.

Additionally, our analyses of the severity of extinction cascades demonstrates that the use of global extinction risk proxies in projections of future ecosystems leads to much more optimistic predictions of biodiversity and functional loss than localised extinction risk proxies. In fact, our local climate-safety-margin simulations yield ecosystem projections similar to our worst-case scenario based on removing the most central species per network. This similarity increases when assuming a worst-case emissions scenario (i.e., SSP585) to define climate-safety margin proxies. Consequently, we find our expectation of localised extinction risk proxies leading to greater ecosystem change than global extinction risk proxies confirmed. Still, we also find a worrying trend of realistic primary extinction sequences leading to similar changes in ecosystems compared to a worst-case-scenario through an ecological network lens.

Consequently, we insist that any projections of future ecosystems take into account localised extinction risk proxies relevant to the scale of assessment and ecological networks rather than easy-to-retrieve global extinction risks. Doing so will minimise the risk of underpredicting change to local and global levels of biodiversity and ecosystem functioning.

### Network-Resilience Landscapes

We find that contemporary approaches to projecting the outcomes of secondary extinctions may be a dangerous oversimplification of biological reality within which species’ dependency on interaction partners and their capability to reallocate resources and interactions vary. We show how the current practice of considering species as secondarily extinct only when all corresponding links are lost with no rewiring taking place (i.e., lower right-hand corner of all network-resilience landscapes shown in this study) may establish potentially overly optimistic biodiversity forecasts (e.g., contrasting number of extant nodes in Figure 1D IV and III). In addition, current practices can also fail to predict novel ecosystem processes established by rewired links (e.g., contrasting number of extant nodes in Figure 1D IV, I and II). Using the network-resilience landscapes, it is possible to explore even more optimistic or far more pessimistic scenarios of future ecosystems quickly and explicitly. Within such an array of ecosystem scenarios, there will be contained realistic outcomes of extinction cascades. We suggest that forecasts of community assemblages and network topology ought to explicitly account for realistic link-loss sensitivity and rewiring probability thresholds.

While our study makes use of the two network resilience mechanisms (link-loss sensitivity and realisation of rewiring potential) as network-wide numerical arguments of the extinction simulation, we believe that both should be incorporated at the species-rather than the network-level to further enhance accuracy of network projections. The NetworkExtinction R package^19^ which we use for our analyses already supports such functionality. We propose that a vital next step in ecological network research will be to focus efforts on quantifying link-loss sensitivity and the realisation potential of novel interactions on a species-by-species basis. Establishing standardised, robust, and accurate protocols of such quantifications enable the use of the network-resilience landscape concept we propose here at even higher degrees of biological realism. This would allow forecasts of ecosystem assemblages and functioning to be augmented drastically effectively arriving at realistic projections rather than optimistic or pessimistic ones as we do here.

### Cascade Directionality

Contrasting the effects of extinction cascade directionality on changes to ecological networks pre-and post-extinction cascade simulation, we find that bottom-up cascades affect networks more adversely than top-down cascades (Figure 5). However, we demonstrate that bidirectional extinction cascades lead to greater change in ecosystem composition and network topology as we hypothesised, particularly due to a larger and arguably more realistic pool of primary extinctions. Therefore, instead of focusing on either cascade direction in isolation as has been done before^31, 32^ we propose that studies of ecological network studies ought to trace extinction cascades of any origin simultaneously as we have done here. In doing so, practitioners may avoid overly optimistic projections of future ecosystems and potentially misidentifying conservation potential and needs.

Our study confirms previous findings of cascade severity on secondary extinctions of the opposing group in mutualistic networks (i.e., primary extinction of plants driving secondary extinction of animals) with bottom-up cascades leading to greater annihilation of the animal species pool than top-down cascades of the plant species pool^18^ (Figure S12). However, these findings hold in our analysis only for the use of global extinction risk proxies, which mirrors the continental extinction risk proxies used in previous research as potentially too coarse to quantify realistic primary extinction risks at the network-level.

### Further considerations

Our study has demonstrated clear avenues for improving ecosystem projections’ accuracy by using information contained in ecological networks. However, we believe that our framework can be enhanced further in future iterations by exploring additional proxies of extinction pressures that have been demonstrated to affect network structure, such as land-use intensity changes^2^. Accounting for changes in species abundances which may induce a shift in network links without necessitating primary extinctions^60^ would be an additional significant advancement for our framework.

Additionally, our study treats climatic niche preferences as static when quantifying climate-safety-margins for each species. However, niche shifts have been proven to be a powerful process of avoiding extinction, even more so than range shifts^61^ which would lead to primary extinctions at network locations, but not global extinction. Incorporating knowledge of probable shifts in species niches into considerations of localised extinction risk calculations may change the overall magnitude of ecosystem change across the network-resilience landscapes we present here but would not affect the network resilience mechanisms we have explored.

Moreover, while our study is one of few which explore the effects of rewiring potential in the face of extinction events, we realise the limitations in our approach. For instance, our approach of allowing only whole link rewiring rather than partial link rewiring (e.g., splitting lost interaction strength across a set of novel or already established partners) may be regarded as overly simplistic. Predicting novel interactions is complex, and experimentation may be needed^62^ rather than deriving rewiring potential from functional trait approaches as we have done. Exploring already demonstrated effects of phylogenetic conservation of interaction preferences^63^ and powerful practices of ecological interaction inference^64, 65^ may help improve the practice of studying rewiring potential in ecological networks.

Finally, our framework suffers from another drawback shared with most contemporary approaches predicting changes in ecological networks – failure to account for introduction of novel species. The approach we present in this study treats node sets for each ecological network as static, only allowing for the removal but not the addition of any new nodes. Previous research has demonstrated that secondary extinctions can be counteracted through colonisation events^66, 67^ (i.e., the introduction of novel species) as long as colonisation is realised quickly while secondary extinctions are not^68^ which mirrors our approach of allowing rewiring before secondary extinction risk is quantified. We believe that closing the knowledge gap of how to account for species introductions into extinction cascade analyses under climate change is particularly timely due to the demonstrated effectiveness of invasion success increasing in warming conditions^69^.

## Conclusion

Ecological networks have proven helpful in rendering more accurate projections of ecosystem change under climate change than single species approaches, mainly through their capability of tracking extinction cascades. Here, we have demonstrated that the assessment of extinction cascade severity is driven by three critical factors which have gone underappreciated in contemporary research practices: (1) risk of primary extinction ought to be quantified using local extinction risk proxies rather than global assessments, (2) incorporating network resilience mechanisms allows the exploration of more realistic extinction cascade outcomes, and (3) bidirectional extinction cascades lead to greater ecosystem level changes than bottom-up or top-down cascades. By implementing these guidelines, we have conclusively shown that current ecosystem forecast practices are likely to underpredict threats to ecosystems. While we point out ways our framework can be enhanced, we are aware that the threat to biodiversity is acute and waiting for development of full-system models might postpone elucidation of conservation potential to a detrimental degree^70^. While we have explored potential ecosystem projections rather than expected projections for each target ecosystem, our study framework represents a powerful approach to exploring more realistic ranges of future ecosystem scenarios than previous approaches. This new perspective will allow more effective targeting of conservation efforts.

## Conflict of Interest

The authors declare no conflicts of interest.

## Authors’ contributions

E.K and A.O conceptualised this study. E.K. created all R scripts necessary for analyses. E.K. carried out the analysis. E.K and A.O interpreted the results. All authors contributed critically to the drafts and gave final approval for publication. This research was supported by the Aarhus University Research Foundation Strat-up Grant (grant no. AUFF-2018-7-8) to AO.

## Data availability

All code required for data retrieval, data handling, extinction simulation, and post-extinction analyses is available at https://github.com/ErikKusch/Ecological-Network-Extinction-Simulations. Fricke 2021 data containing network and functional trait datasets is available at https://zenodo.org/record/5565123#.Yuk_URxByU. We are ready to submit code and data to dryad upon acceptance of the manuscript.

## Supplementary Material

**FIGURE S1.**
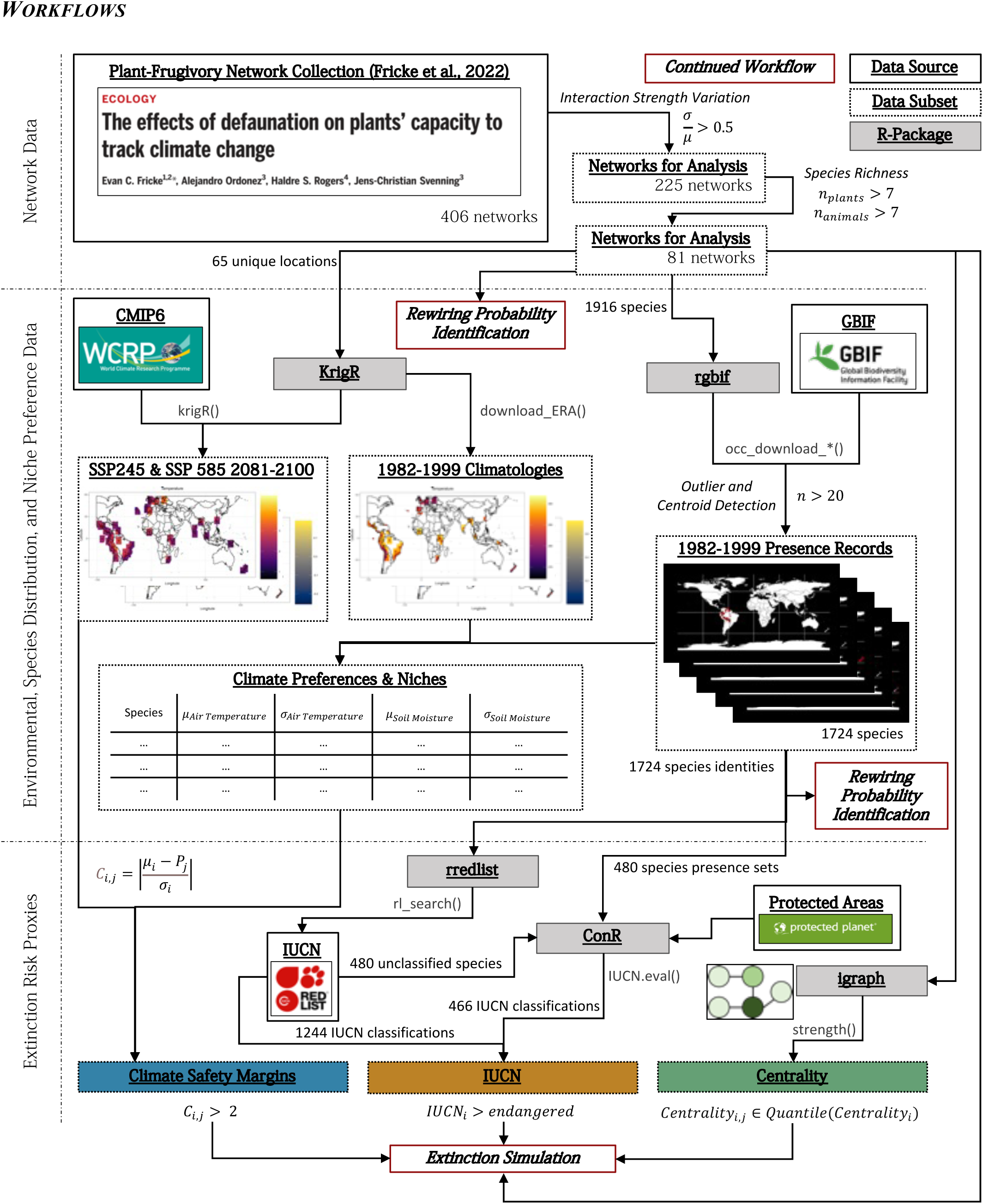
DATA HANDLING & EXTINCTION PROXY IDENTIFICATION. Network data was obtained from Fricke et al, 2022 and subsetted as described in the manuscript. The main text also details how extinction risk proxies and the necessary data for those computations were obtained in a verbose manner.

**FIGURE S2.**
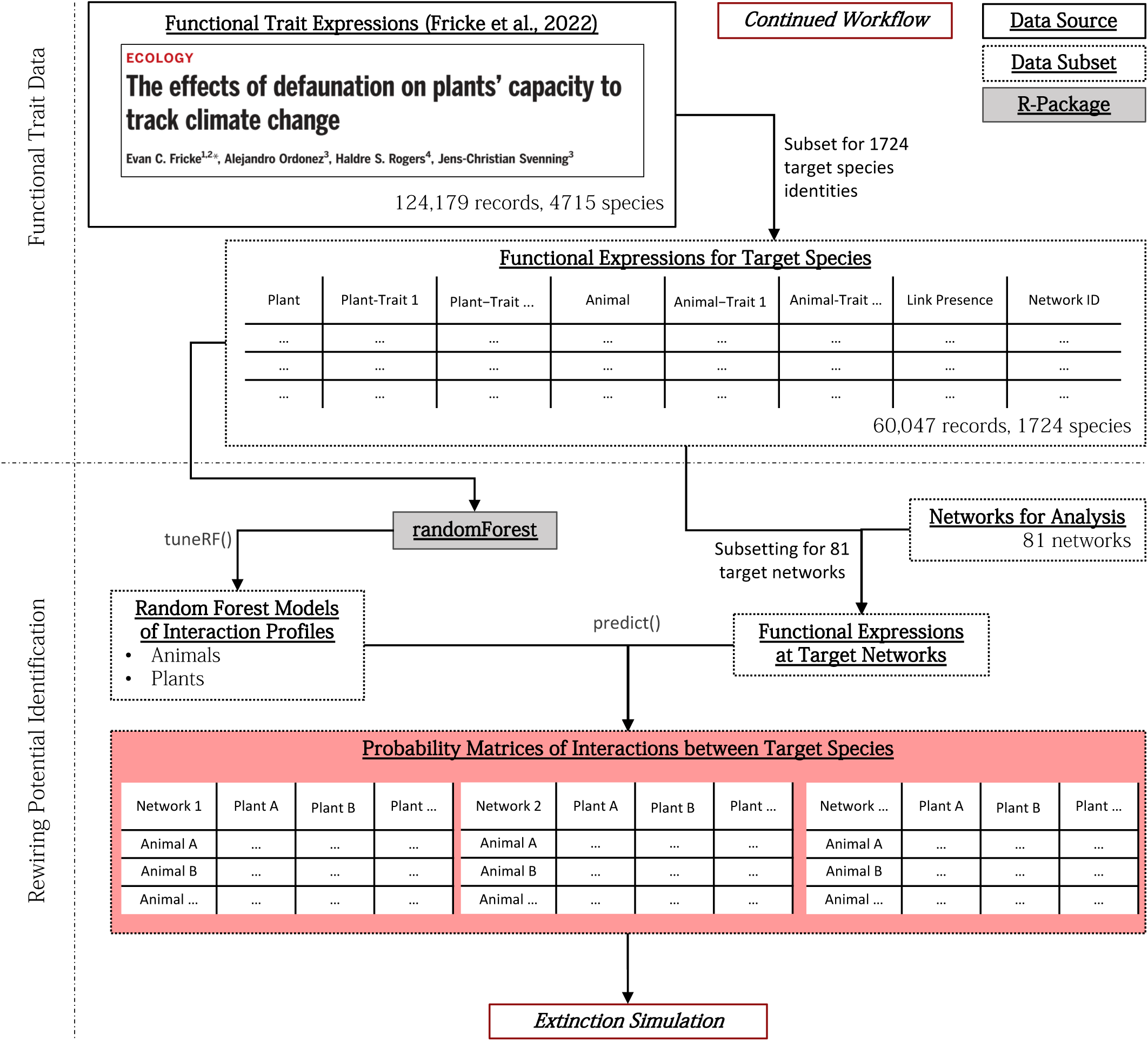
REWIRING PROBABILITY QUANTIFICATION. Rewiring potential was identified by building random forest models on the entire Fricke et al., 2022 data base while predictions of rewiring probability were made on a species-and network-specific basis using corresponding subsets of the data.

**FIGURE S3.**
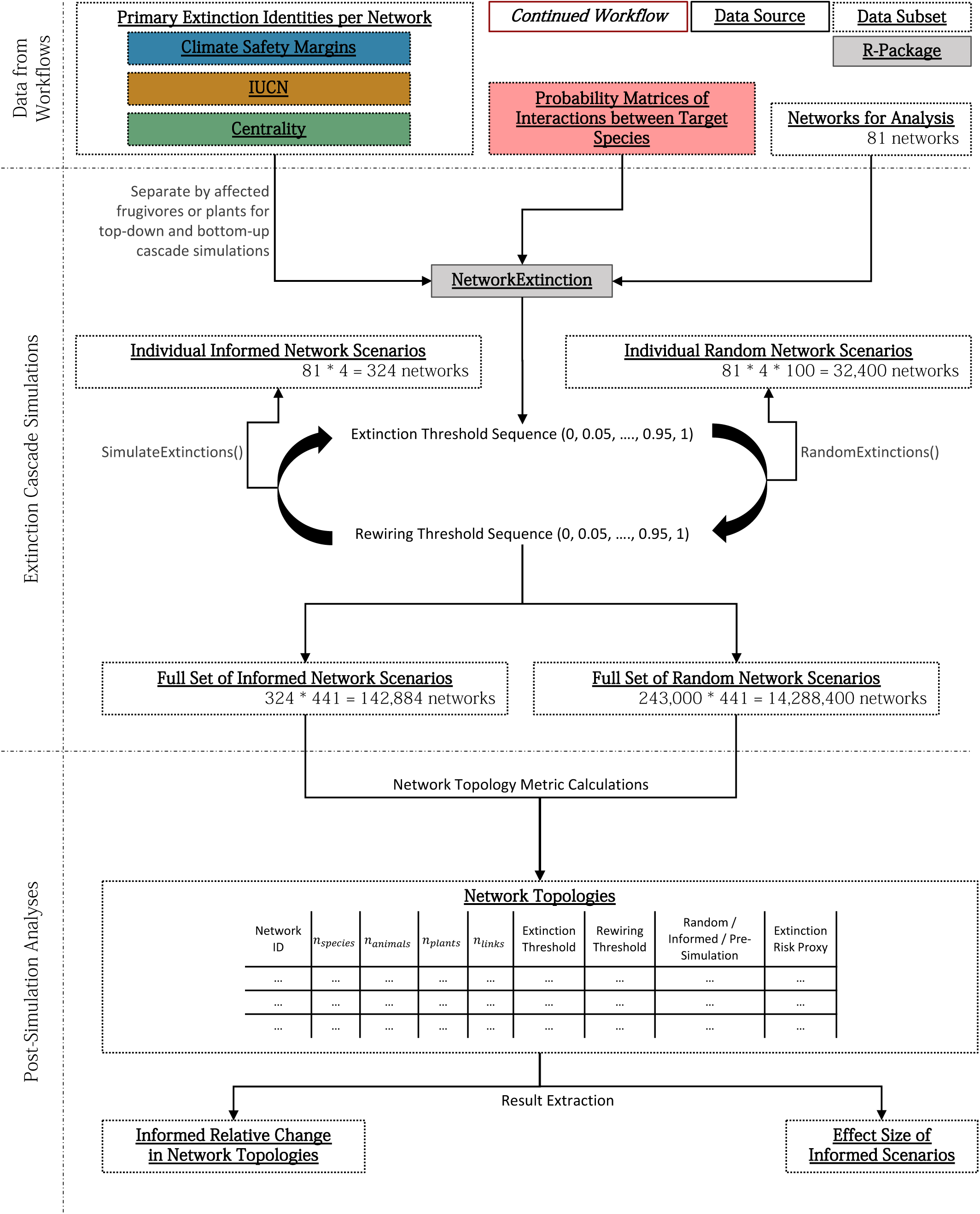
EXTINCTION SIMULATION. Simulations of extinction cascades were executed using functionality of the newly developed NetworkExtinction R package and covered the resilience landscape at 441 distinct locations per network. The extinction simulation procedure was repeated for bottom-up, top-down, and bidirectional extinction cascades.

**FIGURE S4.**
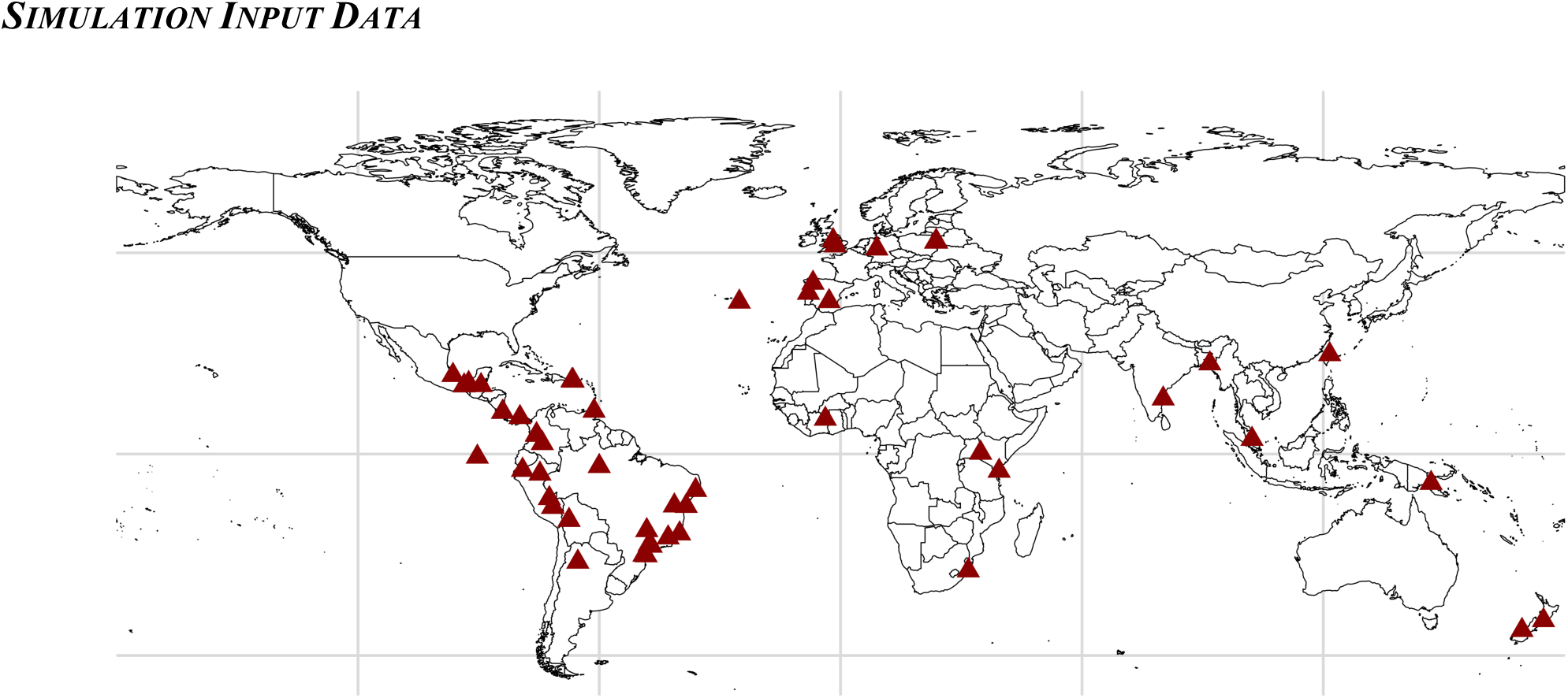
NETWORK OBSERVATION LOCATIONS. Analysis network observations represent a set of global locations.

**FIGURE S5.**
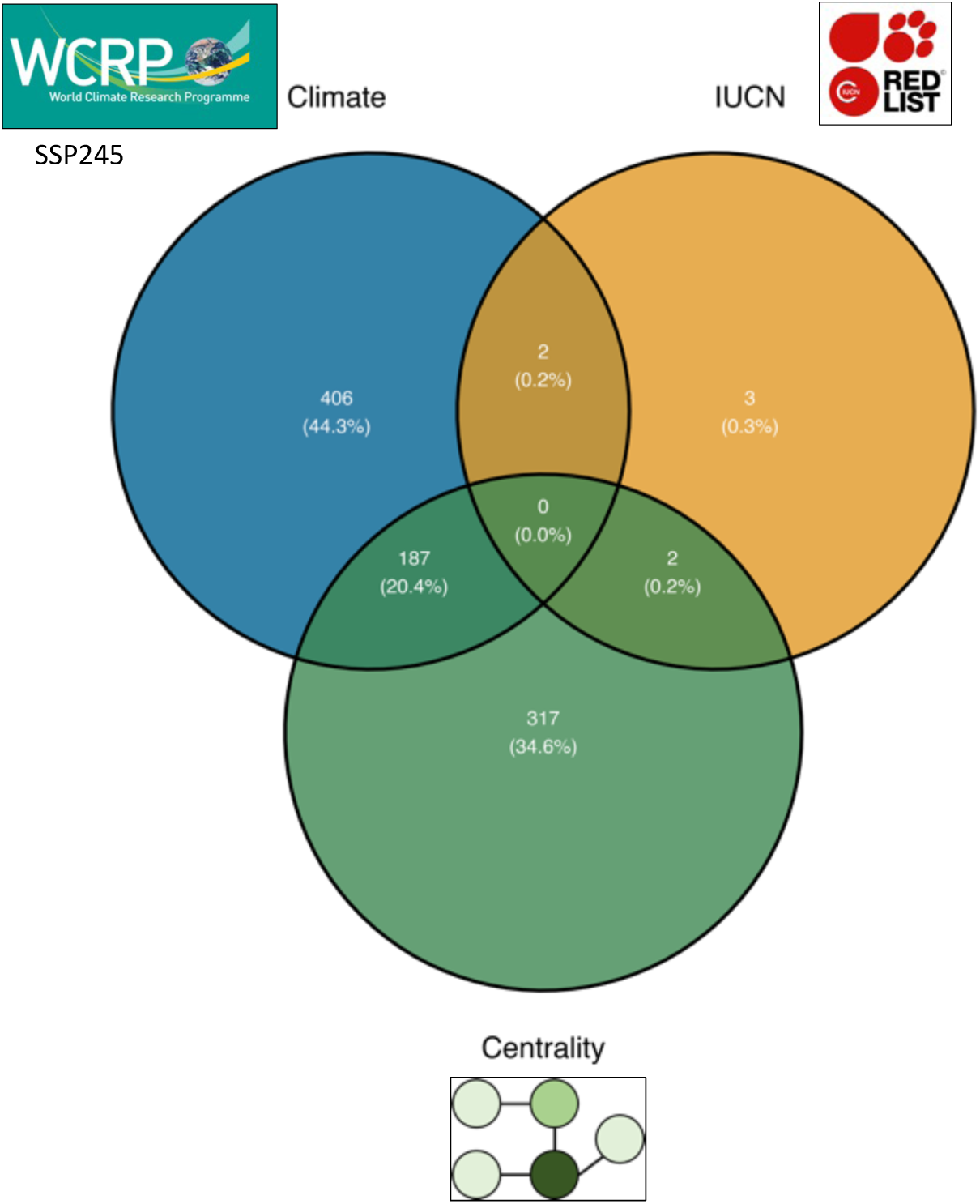
BIDIRECTIONAL PRIMARY EXTINCTION IDENTIFICATION. Allowing animal and plant species to be identified as primary extinction nodes in all analysis networks leads to a far greater number of primary extinctions when using the localised climate-safety-margins and node-centrality proxies as opposed to the global IUCN extinction risk proxy.

**FIGURE S6.**
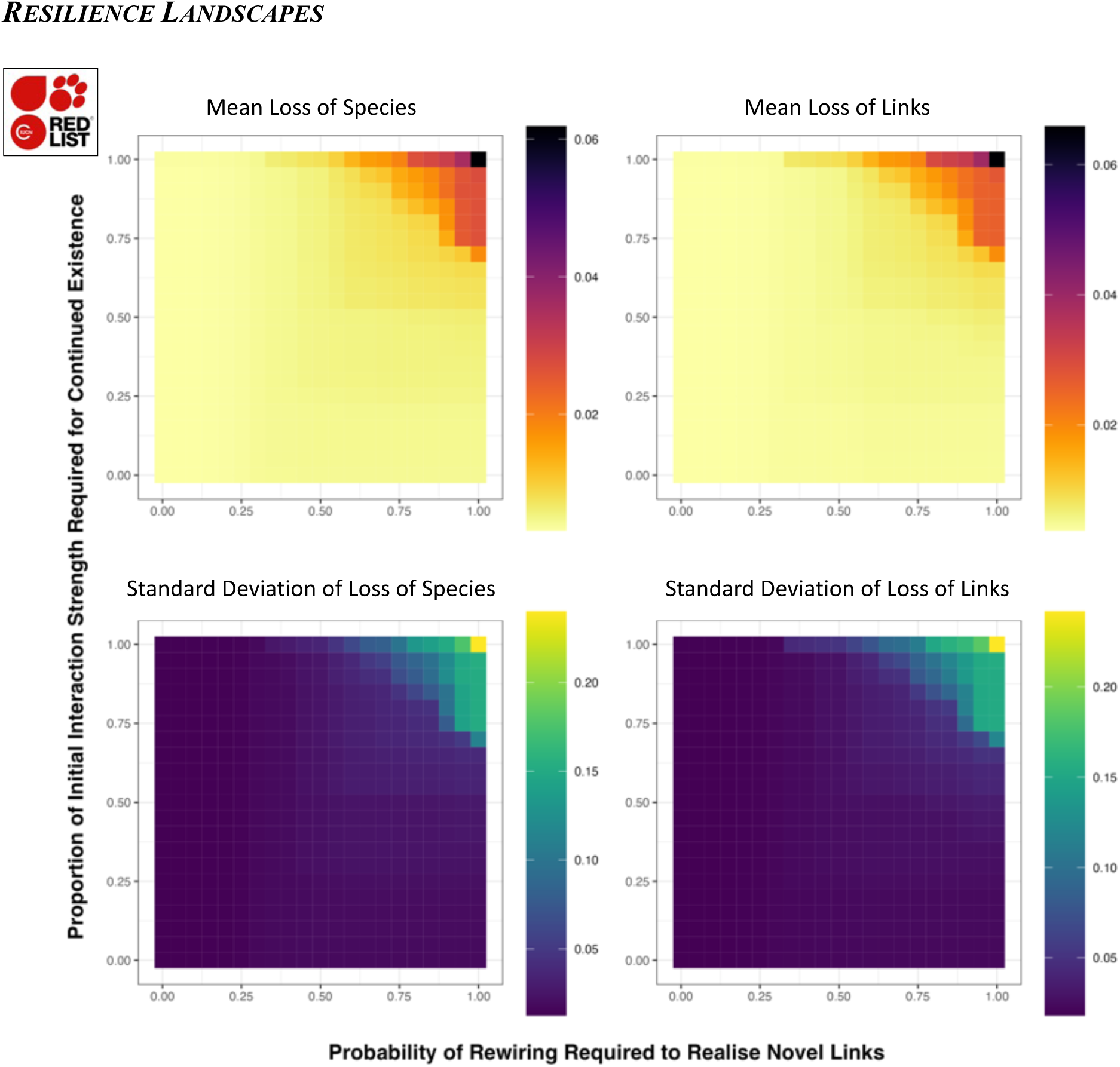
BIDIRECTIONAL IUCN-INFORMED SIMULATION OUTCOMES. These are the means standard deviations simulation outcomes for bidirectional extinction cascades whose primary extinctions were derived from IUCN classifications.

**FIGURE S7.**
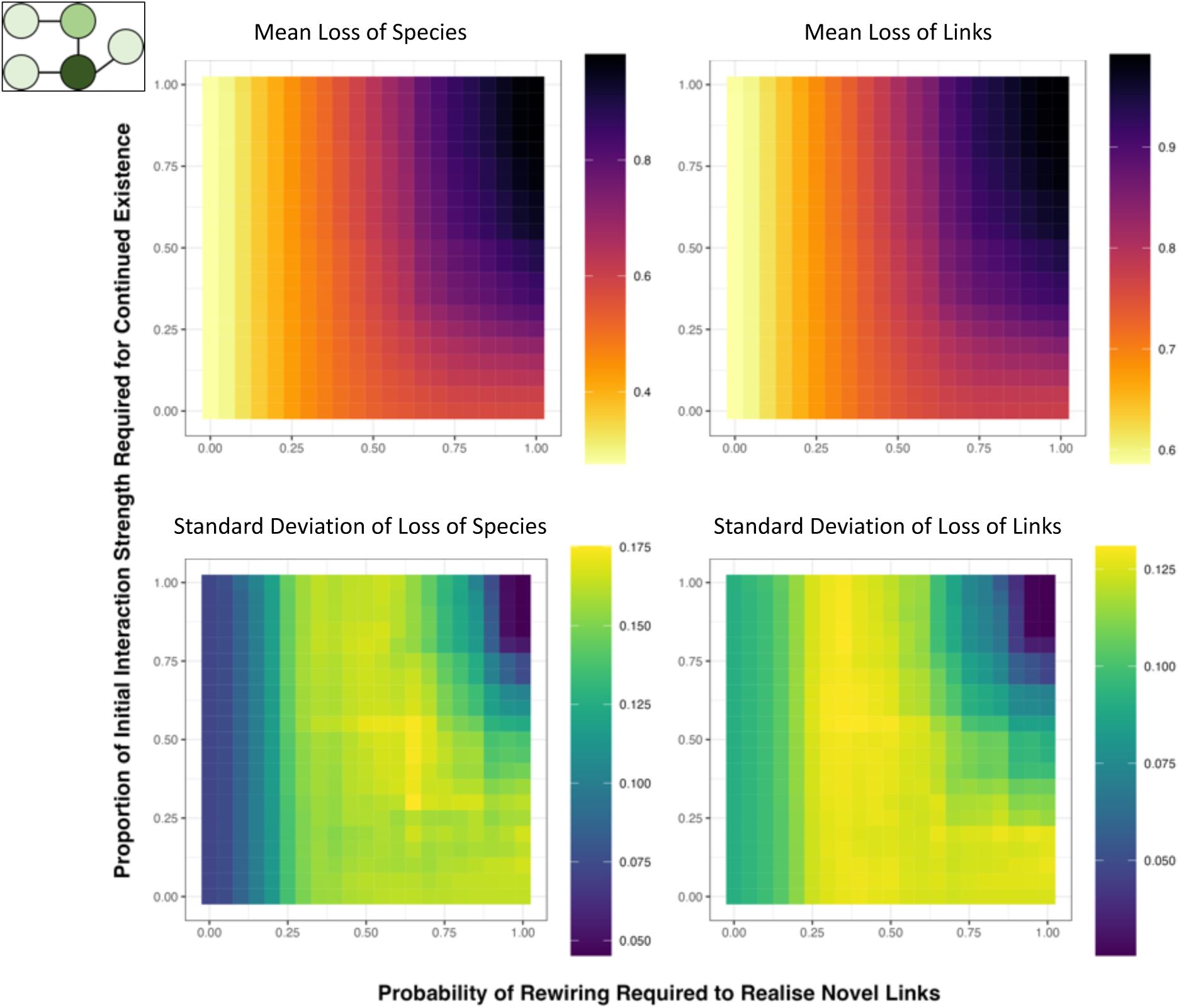
BIDIRECTIONAL CENTRALITY-INFORMED SIMULATION OUTCOMES. These are the means standard deviations simulation outcomes for bidirectional extinction cascades whose primary extinctions were derived from centrality proxies.

**FIGURE S8.**
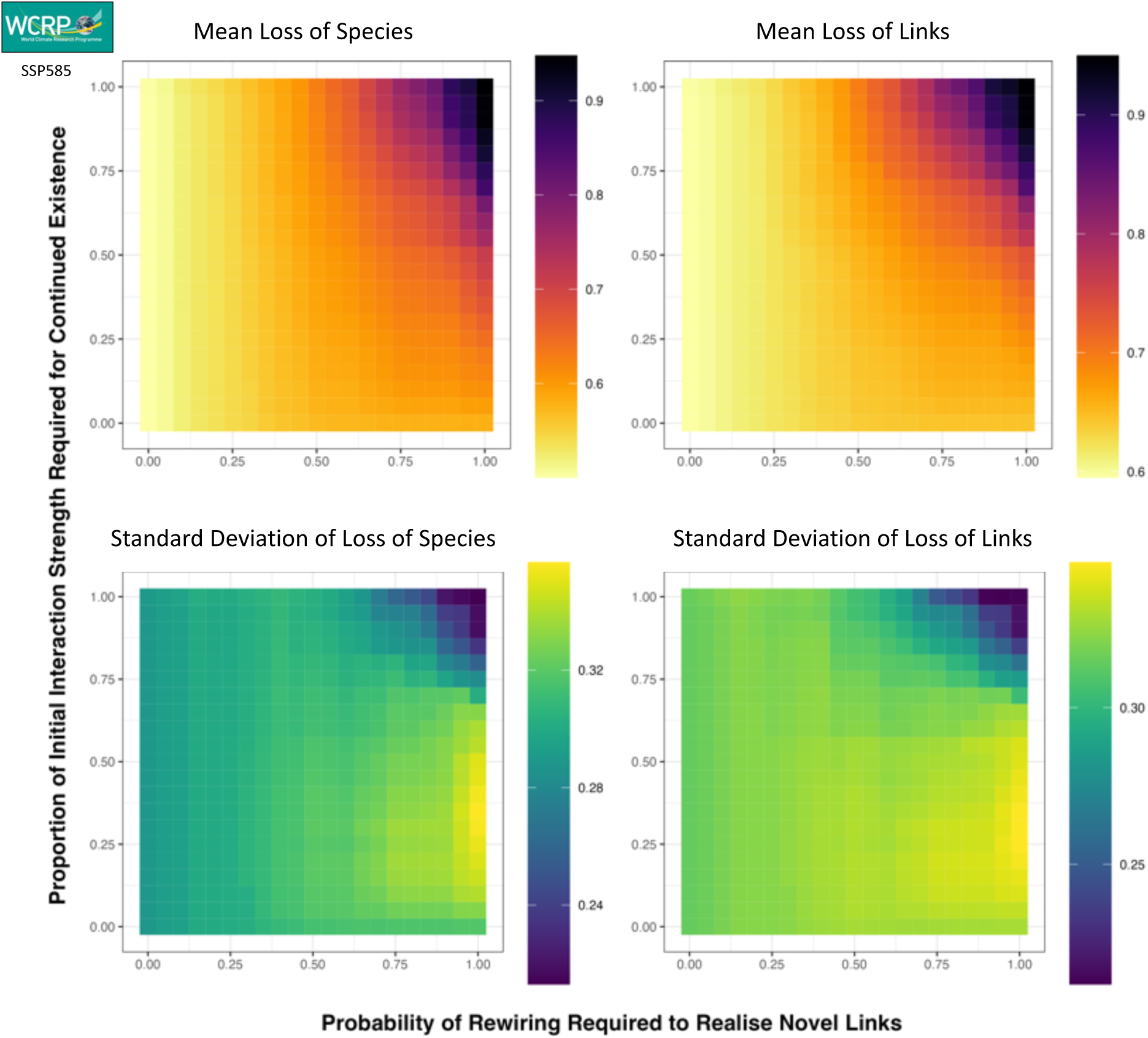
BIDIRECTIONAL CENTRALITY-INFORMED SIMULATION OUTCOMES. These are the means standard deviations simulation outcomes for bidirectional extinction cascades whose primary extinctions were derived from SSP585 climate-safety-margins.

**FIGURE S9.**
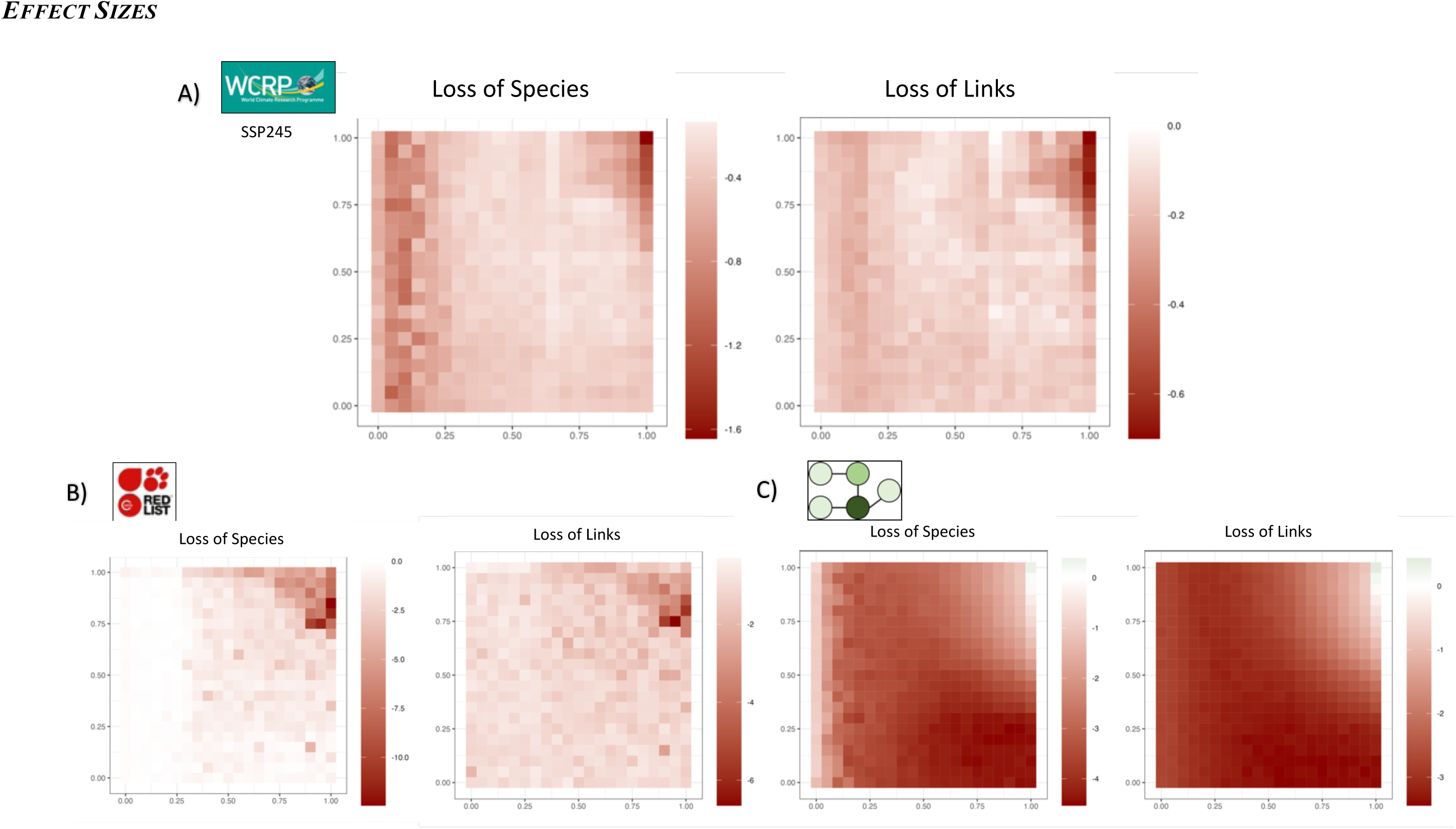
BIDIRECTIONAL EXTINCTION SIMULATION EFFECT SIZES. These are the effect sizes of changes in ecosystem and network metrics reported in Figure 3 in the manuscript and Figures S6 (top-row) and S7 (top-row).

**FIGURE S10.**
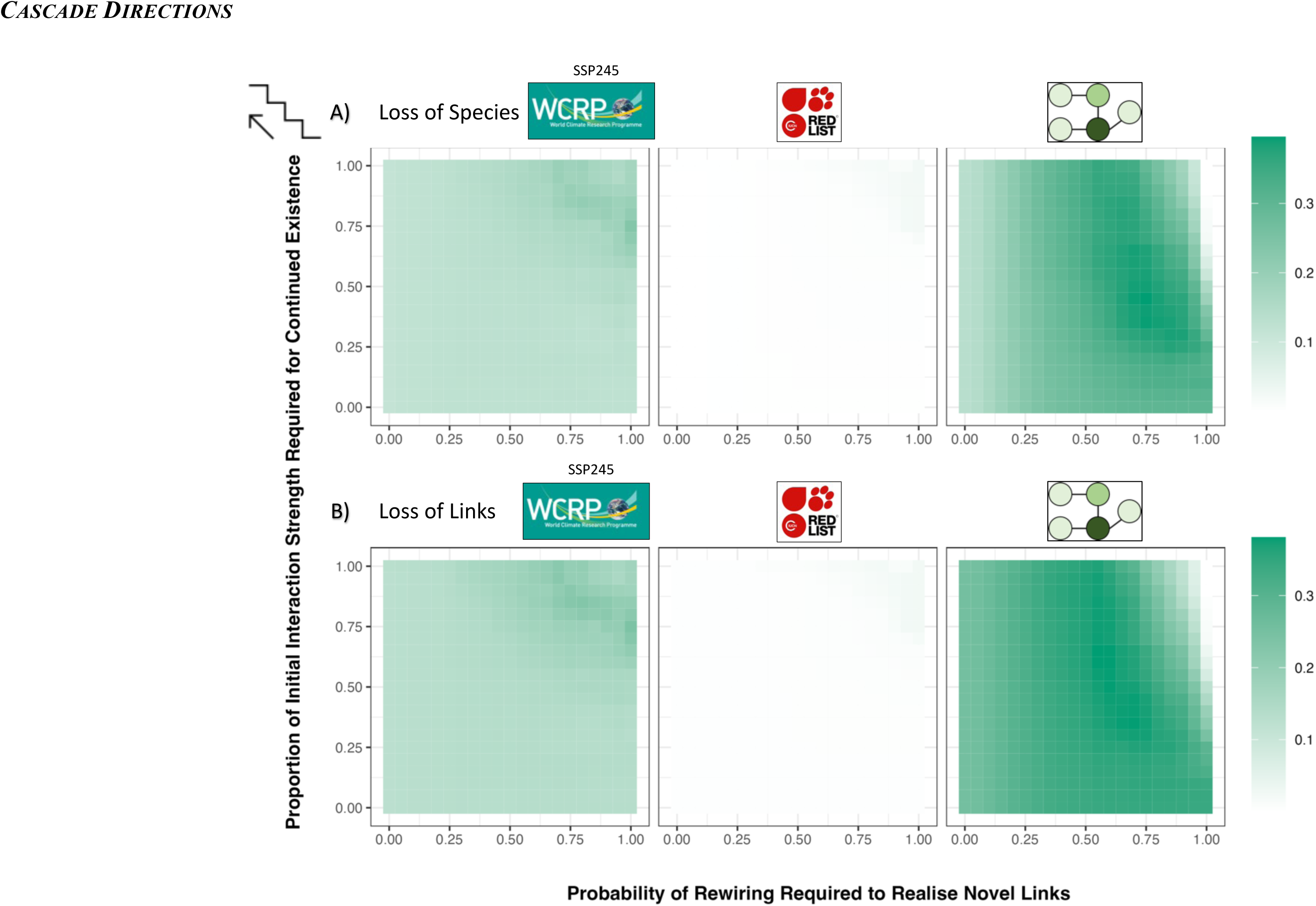
CONTRASTING ECOSYSTEM CHANGE METRICS FOR BIDIRECTIONAL CASDACDES WITH BOTTOM-UP CASCADES. Our analyses demonstrate that bidirectional extinction cascades lead to greater change in ecosystems than bottom-up cascades when using local extinction risk proxies to define primary extinction sequences.

**FIGURE S11.**
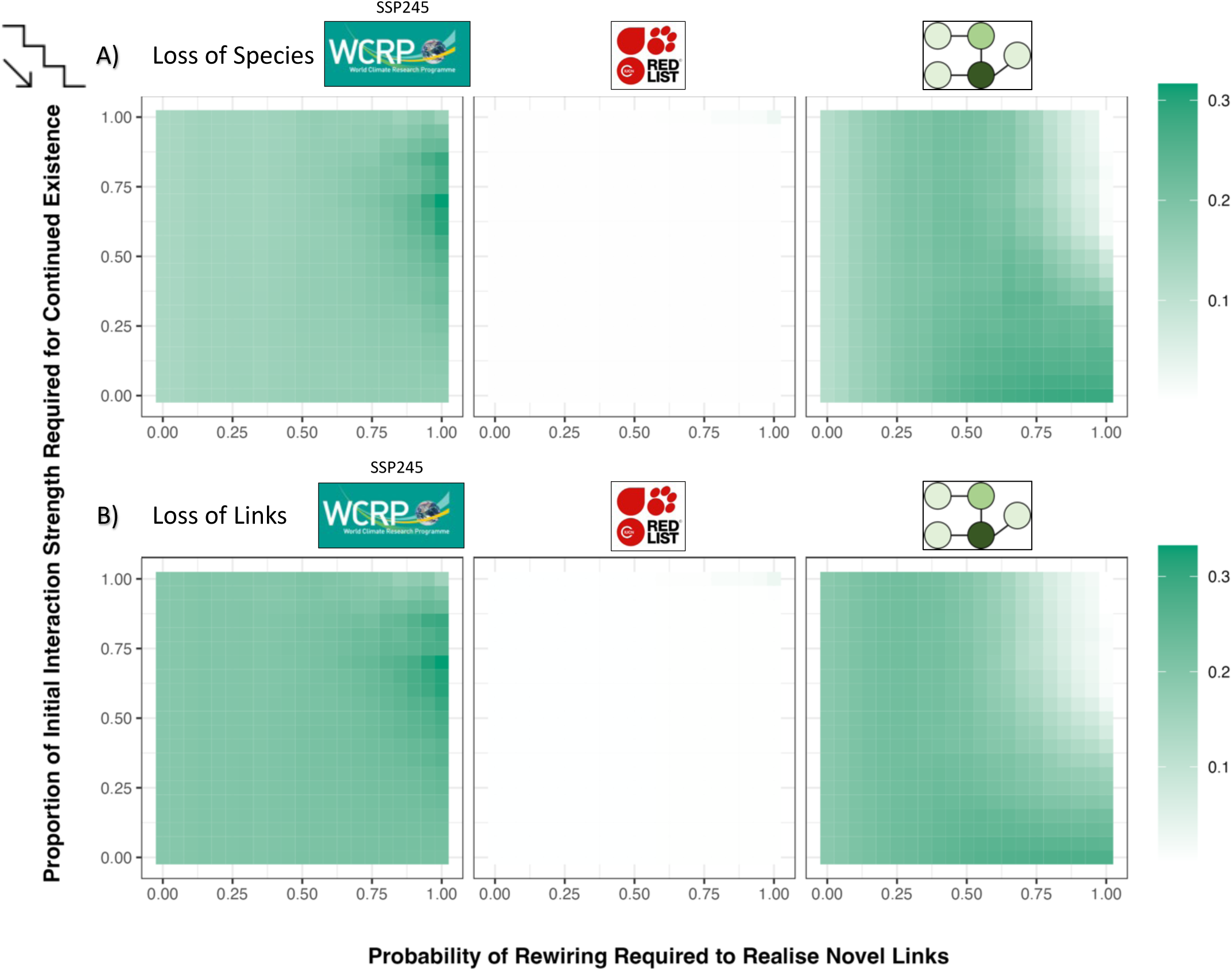
CONTRASTING ECOSYSTEM CHANGE METRICS FOR BIDIRECTIONAL CASDACDES WITH TOP-DOWN CASCADES. Our analyses demonstrate that bidirectional extinction cascades lead to greater change in ecosystems than top-down cascades when using local extinction risk proxies to define primary extinction sequences.

**FIGURE S12.**
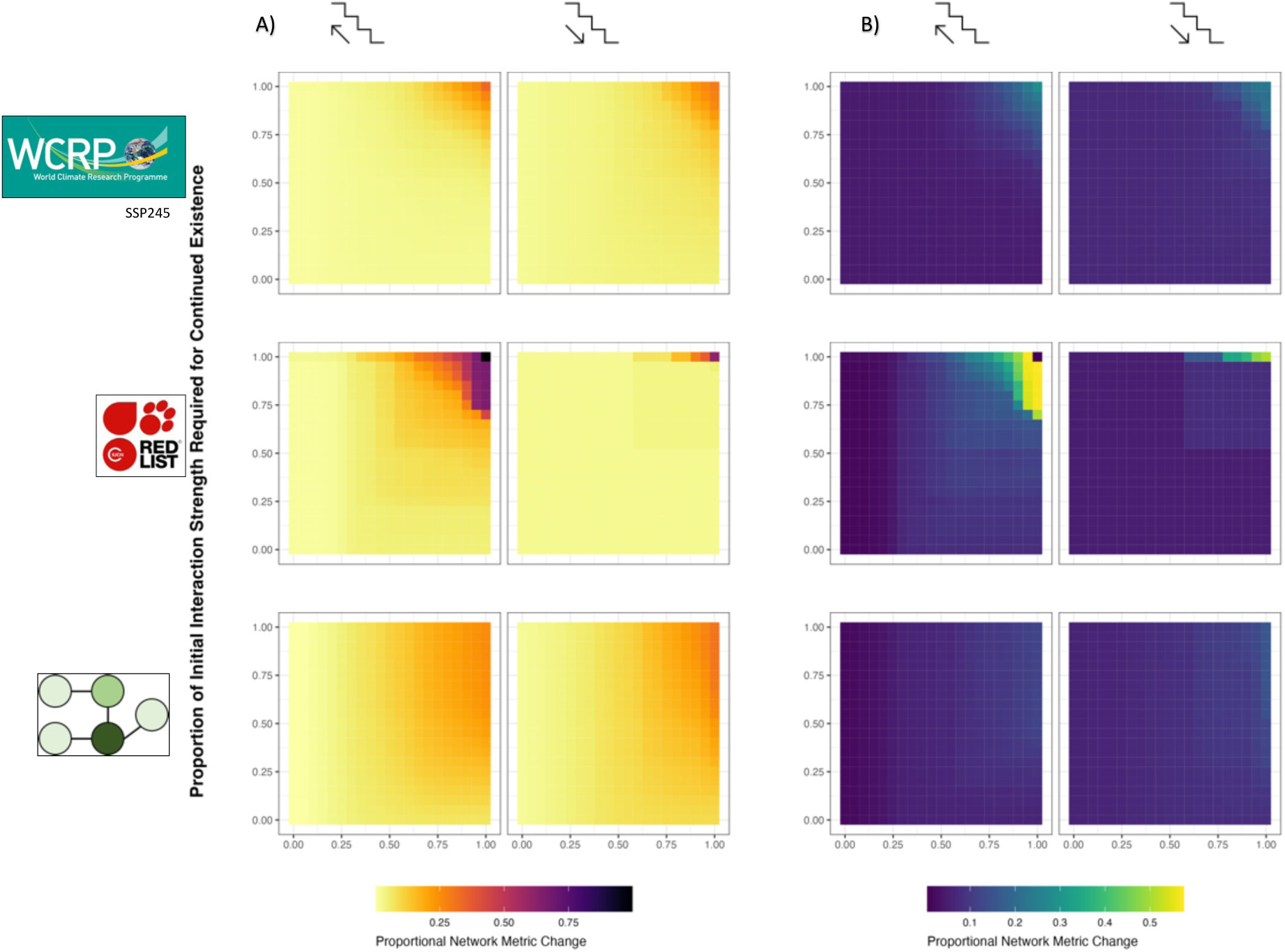
NETWORK SENSITIVITY TO SPECIES-LOSS. Network sensitivity (loss of animal species as compared to loss of plant species and vice versa, Schleuning et al., 2016) confirms previous findings by Schleuning et al., 2016 with bottom-up extinction cascades leading to greater loss of animal species (first column in A) than top-down extinction cascades do for plant species (second column in A). This is particularly striking for IUCN-informed primary extinction sequences, and less so for local extinction risk proxy driven extinction sequences. B) shows the standard deviations of network sensitivity scores across the respective network resilience landscapes shown in A).

## Notes

https://github.com/ErikKusch/Ecological-Network-Extinction-Simulations

### Competing Interest Statement

The authors have declared no competing interest.

### Summary of Updates

Manuscript contained an incorrect reference to a GBIF download that was not used in the analyses. This has been remedied by referencing the DOIs of the GBIF data downloads that were actually used.

## References

1. Barnosky, A. D. et al. Has the Earth’s sixth mass extinction already arrived? Nature vol. 471 51–57 at https://doi.org/10.1038/nature09678 (2011).

2. Felipe-Lucia, M. R. et al. Land-use intensity alters networks between biodiversity, ecosystem functions, and services. Proc. Natl. Acad. Sci. 117, 28140–28149 (2020).

3. Reyer, C. P. O. et al. Forest resilience and tipping points at different spatio-temporal scales: approaches and challenges. J. Ecol. 103, 5–15 (2015).

4. Scheffer, M., Carpenter, S., Foley, J. a, Folke, C. & Walker, B. Catastrophic shifts in ecosystems. Nature 413, 591–596 (2001).

5. Seidl, R., Spies, T. A., Peterson, D. L., Stephens, S. L. & Hicke, J. A. REVIEW: Searching for resilience: addressing the impacts of changing disturbance regimes on forest ecosystem services. J. Appl. Ecol. 53, 120–129 (2016).

6. Schickhoff, U. et al. Do Himalayan treelines respond to recent climate change? An evaluation of sensitivity indicators. Earth Syst. Dyn. 6, 245–265 (2015).

7. IPBRES. IPBES - Global assessment - Full report. (2019).

8. Pimm, S. L. et al. The biodiversity of species and their rates of extinction, distribution, and protection. Science (80-.). 344, (2014).

9. Peterson, A. T. & Soberón, J. Species distribution modeling and ecological niche modeling: Getting the Concepts Right. Nat. a Conserv. 10, 102–107 (2012).

10. Biber, M. F., Voskamp, A., Niamir, A., Hickler, T. & Hof, C. A comparison of macroecological and stacked species distribution models to predict future global terrestrial vertebrate richness. J. Biogeogr. 47, 114–129 (2020).

11. Morueta-Holme, N. et al. A network approach for inferring species associations from co-occurrence data. Ecography (Cop.). 39, 1139–1150 (2016).

12. Stephan, P., Mora, B. B. & Alexander, J. M. Positive species interactions shape species’ range limits. Oikos oik.08146 (2021) doi:10.1111/oik.08146.

13. Bascompte, J. & Jordano, P. Plant-Animal Mutualistic Networks: The Architecture of Biodiversity. Annu. Rev. Ecol. Evol. Syst. 38, 567–593 (2007).

14. D’Amen, M., Mod, H. K., Gotelli, N. J. & Guisan, A. Disentangling biotic interactions, environmental filters, and dispersal limitation as drivers of species co-occurrence. Ecography (Cop.). (2018) doi:10.1111/ecog.03148.

15. Pollock, L. J. et al. Understanding co-occurrence by modelling species simultaneously with a Joint Species Distribution Model (JSDM). Methods Ecol. Evol. 5, 397–406 (2014).

16. Fortin, M., Dale, M. R. T. & Brimacombe, C. Network ecology in dynamic landscapes. Proc. R. Soc. B Biol. Sci. 288, rspb.2020.1889 (2021).

17. Strona, G. & Bradshaw, C. J. A. Co-extinctions annihilate planetary life during extreme environmental change. Sci. Rep. 8, 16724 (2018).

18. Schleuning, M. et al. Ecological networks are more sensitive to plant than to animal extinction under climate change. Nat. Commun. 7, 1–9 (2016).

19. Ávila-Thieme, M. I., et al. NetworkExtinction: An R package to simulate extinction propagation and rewiring potential in ecological networks. Methods Ecol. Evol. 2020.10.17.305391 (2023) doi:10.1111/2041-210X.14126.

20. Stuart, S. N. et al. Status and Trends of Amphibian Declines and Extinctions Worldwide. Science (80-.). 306, 1783–1786 (2004).

21. McWilliams, C., Lurgi, M., Montoya, J. M., Sauve, A. & Montoya, D. The stability of multitrophic communities under habitat loss. Nat. Commun. 10, 1–11 (2019).

22. Bellard, C., Bernery, C. & Leclerc, C. Looming extinctions due to invasive species: Irreversible loss of ecological strategy and evolutionary history. Glob. Chang. Biol. gcb.15771 (2021) doi:10.1111/gcb.15771.

23. BirdLife International. STATE OF THE WORLD’S BIRDS. (2018).

24. Bellard, C., Genovesi, P. & Jeschke, J. M. Global patterns in threats to vertebrates by biological invasions. Proc. R. Soc. B Biol. Sci. 283, (2016).

25. Gallagher, R. V., Allen, S. & Wright, I. J. Safety margins and adaptive capacity of vegetation to climate change. Sci. Rep. 9, 1–11 (2019).

26. Higino, G. T., Windsor, F. M., Banville, F. & Dansereau, G. Mismatch between IUCN range maps and species interactions data illustrated using the Serengeti food web. (2022).

27. Kellermann, V. et al. Upper thermal limits of Drosophila are linked to species distributions and strongly constrained phylogenetically. Proc. Natl. Acad. Sci. 109, 16228–16233 (2012).

28. Hodgson, D., McDonald, J. L. & Hosken, D. J. What do you mean, ‘resilient’? Trends Ecol. Evol. 30, 503–506 (2015).

29. Fründ, J. Dissimilarity of species interaction networks: how to partition rewiring and species turnover components. Ecosphere 12, (2021).

30. Carpentier, C., Barabás, G., Spaak, J. W. & De Laender, F. Reinterpreting the relationship between number of species and number of links connects community structure and stability. *Nat*. Ecol. Evol. (2021) doi:10.1038/s41559-021-01468-2.

31. Estes, J. A. et al. Trophic Downgrading of Planet Earth. Science (80-.). 333, 301–306 (2011).

32. Fricke, E. C., Ordonez, A., Rogers, H. S. & Svenning, J.-C. The effects of defaunation on plants’ capacity to track climate change. Science (80-.). 375, 210–214 (2022).

33. Ordonez, A. & Svenning, J.-C. Geographic patterns in functional diversity deficits are linked to glacial-interglacial climate stability and accessibility. Glob. Ecol. Biogeogr. 24, 826–837 (2015).

34. R Core Team. R: A Language and Environment for Statistical Computing. at (2021).

35. Fricke, E. C. evancf/global-dispersal-change: Primary release of data and code. Zenodo https://zenodo.org/record/5565123#.Yuk_URxByUk (2021) doi:10.5281/zenodo.5565123.

36. Ramos-Robles, M., Andresen, E. & Díaz-Castelazo, C. Temporal changes in the structure of a plant-frugivore network are influenced by bird migration and fruit availability. PeerJ 2016, (2016).

37. Albrecht, J. et al. Logging and forest edges reduce redundancy in plant-frugivore networks in an old-growth European forest. J. Ecol. 101, 990–999 (2013).

38. van de Pol, M. et al. Identifying the best climatic predictors in ecology and evolution. Methods Ecol. Evol. 7, 1246–1257 (2016).

39. Walter, J., Jentsch, A., Beierkuhnlein, C. & Kreyling, J. Ecological stress memory and cross stress tolerance in plants in the face of climate extremes. Environ. Exp. Bot. 94, 3– 8 (2013).

40. Kusch, E., Davy, R. & Seddon, A. W. R. Vegetation-memory effects and their association with vegetation resilience in global drylands. J. Ecol. 110, 1561–1574 (2022).

41. Sabater, J. M. ERA5-Land: A new state-of-the-art Global Land Surface Reanalysis Dataset. Earth Syst. Sci. Data (2021) doi:https://doi.org/10.5194/essd-13-4349-2021.

42. Kusch, E. & Davy, R. KrigR—a tool for downloading and statistically downscaling climate reanalysis data. Environ. Res. Lett. 17, 024005 (2022).

43. Eyring, V. et al. Overview of the Coupled Model Intercomparison Project Phase 6 (CMIP6) experimental design and organization. Geosci. Model Dev. 9, 1937–1958 (2016).

44. Davy, R. & Kusch, E. Reconciling high resolution climate datasets using KrigR. Environ. Res. Lett. 16, 124040 (2021).

45. Hawkins, E., Osborne, T. M., Ho, C. K. & Challinor, A. J. Calibration and bias correction of climate projections for crop modelling: An idealised case study over Europe. Agric. For. Meteorol. 170, 19–31 (2013).

46. Quintana Seguí, P., Ribes, A., Martin, E., Habets, F. & Boé, J. Comparison of three downscaling methods in simulating the impact of climate change on the hydrology of Mediterranean basins. J. Hydrol. 383, 111–124 (2010).

47. Ehret, U., Zehe, E., Wulfmeyer, V., Warrach-Sagi, K. & Liebert, J. Should we apply bias correction to global and regional climate model data? Hydrology and Earth System Sciences vol. 16 3391–3404 at https://doi.org/10.5194/hess-16-3391-2012 (2012).

48. GBIF Occurrence Download. (2023) doi:10.15468/dl.fmujjg.

49. GBIF Occurrence Download. (2023) doi:10.15468/dl.skrwmw.

50. Chamberlain, S. et al. rgbif: Interface to the Global Biodiversity Information Facility API. at (2022).

51. Natural Earth. Natural Earth Data. https://www.naturalearthdata.com/ (2021).

52. Chamberlain, S. rredlist: IUCN Red List Client. at (2020).

53. Dauby, G. et al. ConR : An R package to assist large-scale multispecies preliminary conservation assessments using distribution data. Ecol. Evol. 7, 11292–11303 (2017).

54. UNEP-WCMC and IUCN. Protected Planet: The World Database on Protected Areas (WDPA) and World Database on Other Effective Area-based Conservation Measures (WD-OECM). Cambridge, UK: UNEP-WCMC and IUCN www.protectedplanet.net (2022).

55. Dunne, J. A. & Williams, R. J. Cascading extinctions and community collapse in model food webs. Philos. Trans. R. Soc. B Biol. Sci. 364, 1711–1723 (2009).

56. Farine, D. R. & Carter, G. G. Permutation tests for hypothesis testing with animal social network data: Problems and potential solutions. Methods Ecol. Evol. 13, 144–156 (2022).

57. Liaw, A. & Wiener, M. Classification and Regression by randomForest. R News 2, 18– 22 (2002).

58. Bürkner, P. C. Bayesian Item Response Modeling in R with brms and Stan. J. Stat. Softw. 100, (2021).

59. Arias Arone, E. Dieta y estructura trófica de un ensamblaje de murciélagos en un bosque montano de los andes orientales del centro del Perú. (Universidad Nacional Mayor de San Marcos, 2016).

60. Mougi, A. Adaptive plasticity in activity modes and food web stability. PLoS One 17, e0267444 (2022).

61. Román-Palacios, C. & Wiens, J. J. Recent responses to climate change reveal the drivers of species extinction and survival. Proc. Natl. Acad. Sci. 201913007 (2020) doi:10.1073/pnas.1913007117.

62. Alexander, J. M., Diez, J. M., Hart, S. P. & Levine, J. M. When Climate Reshuffles Competitors: A Call for Experimental Macroecology. Trends Ecol. Evol. 31, 831–841 (2016).

63. Strydom, T. et al. Food web reconstruction through phylogenetic transfer of low-rank network representation. (2021) doi:10.32942/osf.io/y7sdz.

64. Bimler, M. D., Mayfield, M. M., Martyn, T. E. & Stouffer, D. B. Estimating interaction matrices from performance data for diverse systems. (2022) doi:doi.org/10.1101/2022.03.28.486154.

65. Kusch, E., Bimler, M. D., Lutz, J. A. & Ordonez, A. Ecological network inference is not consistent across scales or approaches approaches. (2023) doi:10.1101/2023.07.13.548816.

66. Haddou, Y. et al. Delays in biodiversity responses to land cover change lead to extinction debts and colonization credits among US bird communities Authors.

67. Fernandez, F. A. S. et al. Estimating interaction credit for trophic rewilding in tropical forests. Philos. Trans. R. Soc. B Biol. Sci. 373, 20170435 (2018).

68. Vandvik, V. et al. Biotic rescaling reveals importance of species interactions for variation in biodiversity responses to climate change. Proc. Natl. Acad. Sci. U. S. A. 202003377 (2020) doi:10.1073/pnas.2003377117.

69. Sentis, A., Montoya, J. M. & Lurgi, M. Warming indirectly increases invasion success in food webs. Proc. R. Soc. B Biol. Sci. 288, (2021).

70. Barthlott, W. et al. Terminological and methodological aspects of the mapping and analysis of the global biodiversity. Acta Bot. Fenn. 103–110 (1999).

